# Excitatory neuron-specific suppression of the integrated stress response pathway contributes to autism-related phenotypes in a mouse model of fragile X syndrome

**DOI:** 10.1101/2023.04.24.538123

**Authors:** Mehdi Hooshmandi, Vijendra Sharma, Carolina Thörn Perez, Rapita Sood, Konstanze Simbriger, Calvin Wong, Kevin C. Lister, Alba Ureña Guzmán, Trevor D. Bartley, Cecilia Rocha, Gilles Maussion, Emma Nadler, Patricia Margarita Roque, Ilse Gantois, Jelena Popic, Maxime Lévesque, Randal J. Kaufman, Massimo Avoli, Elisenda Sanz, Karim Nader, Randi Jenssen Hagerman, Thomas M. Durcan, Mauro Costa-Mattioli, Jean-Claude Lacaille, Veronica Martinez-Cerdeno, Jay R. Gibson, Kimberly Huber, Nahum Sonenberg, Christos G. Gkogkas, Arkady Khoutorsky

**Affiliations:** Department of Anesthesia and Faculty of Dental Medicine and Oral Health Sciences, McGill University, Montréal, Québec, Canada; Department of Biochemistry, McGill University, Montréal, Québec, Canada; Department of Pharmacology, Medical University Innsbruck, 6020, Innsbruck, Austria; Department of Pathology and Laboratory Medicine, UC Davis School of Medicine; Institute for Pediatric Regenerative Medicine and Shriners Hospitals for Children of Northern California; MIND Institute, UC Davis Medical Center, Sacramento, CA, USA; The Neuro’s Early Drug Discovery Unit (EDDU), Department of Neurology and Neurosurgery, Montreal Neurological Institute-Hospital, McGill University, Montréal, Québec, Canada; Montreal Neurological Institute and Hospital, Departments of Neurology & Neurosurgery and of Physiology, McGill University, 3801 University Street, Montréal, H3A 2B4, Québec, Canada; Degenerative Diseases Program, Sanford-Burnham-Prebys Medical Discovery Institute, La Jolla, California 92037, USA; Department of Cell Biology, Physiology and Immunology, and Neuroscience Institute. Universitat Autònoma de Barcelona, Bellaterra, Spain; Department of Psychology, Faculty of Science, McGill University; Montreal, Canada; MIND Institute and Department of Pediatrics, University of California at Davis Medical Center, Sacramento, CA, USA; Department of Neuroscience, Baylor College of Medicine, Houston, Texas, 77098, USA; Department of Neurosciences, Center for Interdisciplinary Research on Brain and Learning, and Research Group on Neural Signaling and Circuitry, Université de Montréal, Montréal, Québec, Canada; University of Texas Southwestern Medical Center at Dallas, Department of Neuroscience, Dallas, TX 75390-9111, USA; Biomedical Research Institute, Foundation for Research and Technology-Hellas, University Campus, 45110 Ioannina, Greece; Alan Edwards Centre for Research on Pain, McGill University, Montréal, Québec, Canada

**Keywords:** mRNA translation, integrated stress response, autism, fragile X syndrome

## Abstract

Dysregulation of protein synthesis is one of the key mechanisms underlying autism spectrum disorder (ASD). However, the role of a major pathway controlling protein synthesis, the integrated stress response (ISR), in ASD remains poorly understood. Here, we demonstrate that the main arm of the ISR, eIF2α phosphorylation (p-eIF2α), is suppressed in excitatory but not inhibitory neurons in a mouse model of fragile X syndrome (FXS; *Fmr1*^−/y^). We further show that the decrease in p-eIF2α is mediated via activation of the mTORC1. Genetic reduction of p-eIF2α only in excitatory neurons is sufficient to increase general protein synthesis and cause autism-like behavior. In *Fmr1*^−/y^ mice, genetic restoration of p-eIF2α solely in excitatory neurons reverses elevated protein synthesis and rescues autism-related phenotypes. Thus, we reveal a previously unknown causal relationship between excitatory neuron-specific translational control via the ISR pathway, general protein synthesis and core phenotypes reminiscent of autism in a mouse model of FXS.

## Introduction

Dysregulation of protein synthesis is thought to underlie autism-related phenotypes in fragile X syndrome (FXS)^1^ and several other neurodevelopmental disorders.^2–6^ Increased general protein synthesis in FXS, which has been reported in humans^7, 8^ and animal models,^9, 10^ is believed to critically contribute to FXS pathophysiology. The mechanism driving the robust stimulation of global protein synthesis in FXS is not well understood. Several studies have investigated cell-type-specific changes in mRNA translation,^11–14^ yet it is unclear whether elevated global protein synthesis in FXS is observed in all brain cell types. This is highly relevant for autism spectrum disorders, as single-cell genomics of patient cortex revealed that excitatory neurons are amongst the main cell types preferentially affected in autism.^15^ Previous studies have largely focused on the upregulation of the mechanistic/mammalian target of rapamycin complex 1 (mTORC1) pathway as the main driver of elevated protein synthesis and distinct behavioral deficits in animal models of FXS.^16–19^ However, the role of a cardinal signaling pathway controlling general protein synthesis, the integrated stress response (ISR),^20, 21^ in mediating autism-related phenotypes in *Fmr1*^−/y^ mice, a commonly used FXS model, is hitherto unknown. The ISR is a highly conserved cellular mechanism that suppresses general protein synthesis during stress via phosphorylation of the α-subunit of eukaryotic translation initiation factor 2 (p-eIF2α).^20, 22, 23^ Conversely, a decrease in p-eIF2α stimulates general protein synthesis. Here, we show that the ISR is suppressed in the brain of *Fmr1*^−/y^ mice. Notably, we reveal that the p-eIF2α is reduced only in excitatory but not inhibitory neurons. The decrease in p-eIF2α in *Fmr1*^−/y^ mice is mediated via excitatory neuron-specific upregulation of the mTORC1 activity and is accompanied by an elevation of general protein synthesis in excitatory neurons. Using mouse genetics, we show that low levels of p-eIF2α in excitatory neurons, but not all cell types, are sufficient to cause core phenotypes reminiscent of autism. In *Fmr1*^−/y^, reduced eIF2α phosphorylation in excitatory neurons substantially contributes to molecular and behavioral deficits as normalization of p-eIF2α rescued elevated protein synthesis, exaggerated long-term depression (LTD), social behavior deficits, and alleviated audiogenic seizures. Thus, suppression of the ISR in excitatory neurons plays an important role in mediating autism-related phenotypes in a mouse model of FXS.

## Results

### p-eIF2α is decreased in *Fmr1*^−/y^ excitatory neurons

To assess the activity of the ISR pathway in *Fmr1*^−/y^ mice, we measured p-eIF2α levels in tissue lysates from the hippocampus, cortex, amygdala, striatum, and cerebellum, which are key brain areas implicated in autism spectrum disorders (ASD).^24, 25^ We found a significant decrease in p-eIF2α in *Fmr1*^−/y^ mice in the cortex, hippocampus, and amygdala, but not in the striatum and cerebellum (decrease in p-eIF2α; hippocampus, 44.83%, Figure 1A; cortex, 52.41%, Figure 1B; amygdala, 23.3%, Figure 1C; striatum, Figure S1A; cerebellum, Figure S1B). Examination of p-eIF2α during postnatal brain development showed that the differences between wild-type and *Fmr1*^−/y^ mice emerged during postnatal weeks 5-8 (hippocampus, Figure S1C; cortex, Figure S1D; and amygdala, Figure S1E). To determine whether the ISR is also dysregulated in the brain of individuals with FXS, we assessed p-eIF2α in postmortem human brain tissue, which revealed a trend toward a decrease in p-eIF2α in individuals with FXS compared with their matching controls in two cortical areas (Brodmann area 22, n = 7 for control and n = 6 for FXS, *p* = 0.0989, Figure S1F; and Brodmann area 46, n = 7 for control and n = 5 FXS, *p* = 0.1796, Figure S1G; nested analysis of both areas, *p* = 0.0238, Figure S1H, table S1 includes information on human samples). In *Fmr1*^−/y^ mice, quantitative immunohistochemical (IHC) analysis showed that p-eIF2α is decreased in excitatory neurons (labeled with Ca^2+^-calmodulin-dependent protein kinase 2 (CaMK2α)) (hippocampus, Figure 1D; cortex, Figure 1E; amygdala, Figure S2A) but not inhibitory neurons (labeled with glutamic-acid decarboxylase 67 (GAD67)) (hippocampus, Figure 1F; cortex, Figure 1G; amygdala, Figure S2B). No difference was found in total eIF2α protein levels in excitatory or inhibitory neurons (Figures S2C-H). Phosphorylation of eIF2α decreases the availability of the translation preinitiation complex and inhibits translation initiation.^20^ Conversely, the reduction of p-eIF2α stimulates general translation. To study whether the decrease in p-eIF2α in excitatory neurons is accompanied by an increase in translation, we assessed general protein synthesis using azidohomoalanine (AHA)-based metabolic labeling (FUNCAT)^26^ (Figure 1H). Mice were injected with AHA intraperitoneally, and AHA incorporation into nascent polypeptides in the brain was assessed 3 hrs later (Figure 1I). The validity of this approach was confirmed by showing that the protein synthesis inhibitor anisomycin blocks AHA incorporation (Figure S2I). In these (and subsequent) experiments, we focused on two central brain areas linked to autism, the hippocampus and cortex. Consistent with the decrease in p-eIF2α, general protein synthesis was increased in excitatory neurons (increase in AHA incorporation; hippocampus, 63.77%, Figure 1J; cortex, 65.25%, Figure 1K) but not inhibitory neurons (hippocampus, Figure 1L; cortex, Figure 1M) in *Fmr1*^−/y^ mice compared with wild-type animals.

**Figure 1.**
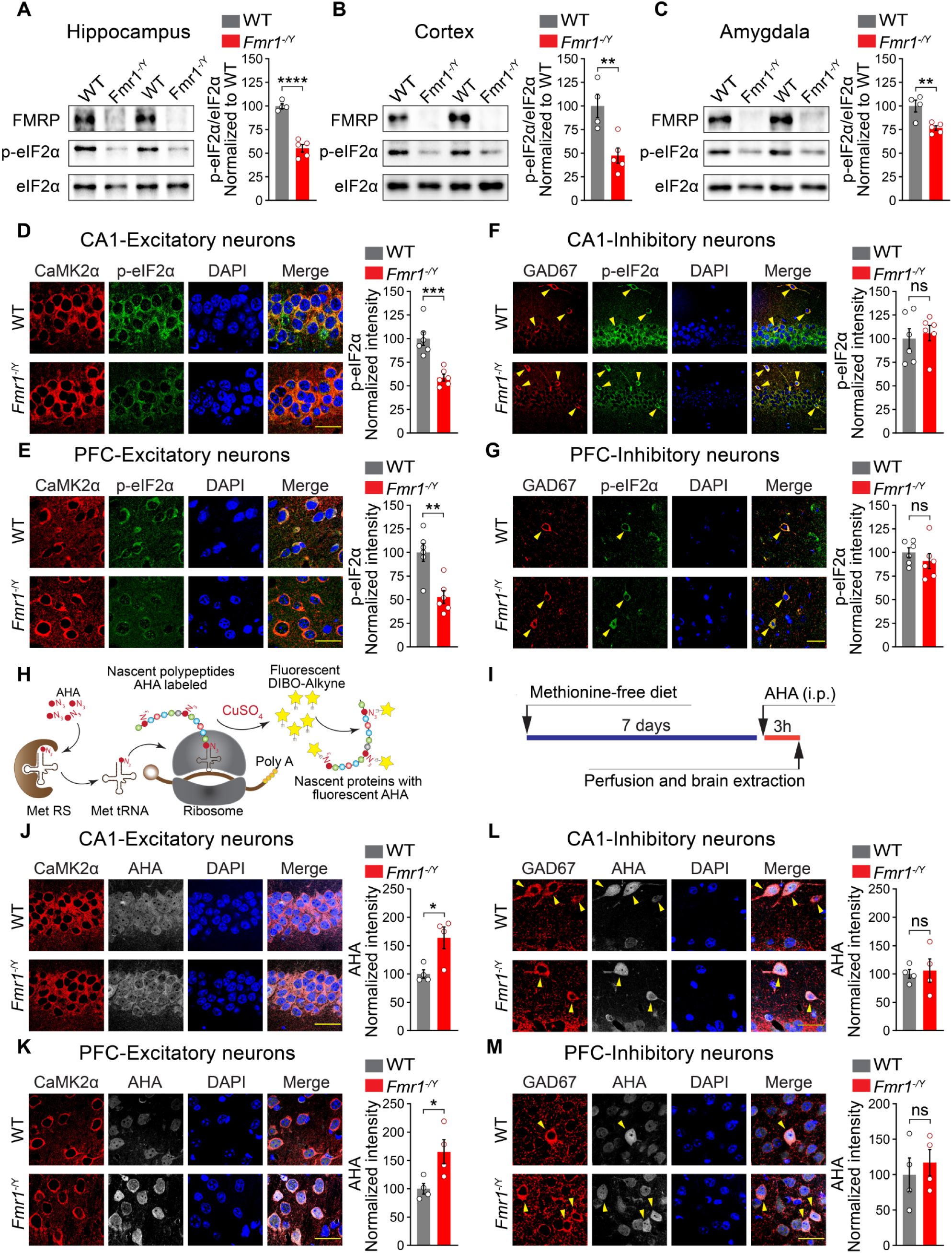
*Fmr1*^-/y^ mice show a reduction in p-eIF2α and an increase in global protein synthesis in excitatory but not inhibitory neurons. (A-C) Representative immunoblotting (left) and quantification (right) showing reduced p-eIF2α in *Fmr1*^-/y^ mice (n = 5) compared with WT mice (n = 4) in hippocampus (A, WT versus *Fmr1*^-/y^, *t* = 8.818, *p* < 0.0001), cortex (B, WT versus *Fmr1*^-/y^, *t* = 3.681, *p* = 0.0078), and amygdala (C, WT versus *Fmr1*^-/y^, *t* = 3.790, *p* = 0.0068). (D-G) Immunofluorescent labelling against p-eIF2α (green) in excitatory (CaMK2α-positive, red) and inhibitory (GAD67-positive, red) neurons reveals reduced p-eIF2α in *Fmr1*^-/y^ (n = 6) compared with WT animals (n = 6) in excitatory neurons in CA1 (D, WT versus *Fmr1*^-/y^, *t* = 5.119, *p* < 0.0005) and prefrontal cortex (PFC) (E, WT versus *Fmr1*^-/y^, *t* = 4.084, *p* < 0.0022), but not inhibitory neurons in CA1 (F, WT versus *Fmr1*^-/y^, *t* = 0.444, *p* > 0.05) and prefrontal cortex (G, WT versus *Fmr1*^-/y^, *t* = 0.967, *p* > 0.05). (H) Schematic illustration of the fluorescence non-canonical amino acid tagging (FUNCAT) and (I) experimental design. AHA incorporation (grey), indicating the level of the nascent protein synthesis, is significantly higher in excitatory (CaMK2α-positive, red) neurons in CA1 (J, WT versus *Fmr1*^-/y^*, t* = 3.083, *p* = 0.0216) and PFC (K, WT versus *Fmr1*^-/y^*, t* = 2.78, *p* = 0.032) of *Fmr1*^-/y^ (n = 4) compared with WT (n = 4) mice. No significant differences were found in AHA incorporation in inhibitory (GAD67-positive, red) neurons in CA1 (L, WT versus *Fmr1*^-/y^*, t*(6) = 0.283, *p* > 0.05) and PFC (M, WT versus *Fmr1*^-/y^*, t* = 0.567, *p* > 0.05) of *Fmr1*^-/y^ (n = 4) compared with WT (n = 4) mice. Yellow arrows mark inhibitory neurons. Each data point represents an individual animal. All data are presented as mean ± s.e.m. \**p* < 0.05, \*\**p* < 0.01, \*\*\**p* < 0.001, \*\*\*\**p* < 0.0001, and ns, not significant. Student’s t-test was performed for all the experiments. Scale bars, 25 µm. See also Figures S1, S2, and S3.

Dephosphorylation of eIF2α causes an increase in general protein synthesis but paradoxically represses translation of a subset of mRNAs, many of which are involved in the cellular stress response and harbor upstream open reading frame (uORF) in their 5′ untranslated region (5′ UTR).^27^ Given the low level of p-eIF2α in excitatory neurons of *Fmr1*^−/y^ mice, we hypothesized that translation of mRNAs containing 5′ UTR uORFs should be reduced in this cell type. Previous studies have examined the translational landscape in excitatory neurons of *Fmr1*^−/y^ mice using excitatory neuron-specific translating ribosome affinity purification (TRAP).^28^ Analysis of the published datasets^11, 12^ revealed that mRNAs containing uORFs in their 5′ UTR were substantially enriched in the subset of genes downregulated in excitatory neurons of *Fmr1*^−/y^ mice compared with control animals (Figure S2J-M, *p* < 0.0001, Fisher’s exact test). Thus, consistent with the reduction of p-eIF2α in excitatory neurons of *Fmr1*^−/y^ mice, general protein synthesis is increased and translation of mRNAs containing uORFs is decreased in excitatory neurons.

### Hyperactivated mTORC1 downregulates p-eIF2α

We next investigated the mechanism underlying the reduction of p-eIF2α in *Fmr1*^−/y^ mice. In the brain, the α subunit of eIF2 can be phosphorylated by stress-activated kinases GCN2, PERK, and PKR. eIF2 can also be dephosphorylated by protein phosphatase 1 (PP1) which is recruited to eIF2α by its two regulatory proteins, a constitutively expressed CReP and a stress-induced GADD34.^20, 21, 29^ Western blot analysis of lysates from hippocampus and cortex of *Fmr1*^−/y^ mice showed an increase in GCN2 phosphorylation on Thr899, indicating its activation,^30^ (hippocampus, 54.2%; cortex, 61.3%, Figures S3A, S3B, S3G, and S3H), and no change in the phosphorylation status or expression levels of PERK, and PKR (Figures S3C-F, S3I-L). GADD34, CReP and PP1 protein expression was not altered in *Fmr1*^−/y^ hippocampus and cortex (Figures S3M-P). These results indicate that the activities of eIF2α kinases and phosphatases cannot explain the reduction in eIF2α phosphorylation in the brain of *Fmr1*^−/y^ mice, suggesting the involvement of another mechanism.

mTORC1 is hyperactivated in the brain of *Fmr1*^−/y^ mice.^16, 17, 19^ Furthermore, a study performed in non-neuronal cultures suggested that activation of mTORC1 can lead to eIF2α dephosphorylation,^31^ though the precise molecular mechanism underlying this effect remains unknown. To investigate whether an increase in mTORC1 activity in the brain causes a decrease in p-eIF2α, we administered wild-type mice with a specific brain-penetrant mTORC1 activator, NV-5138.^32^ As expected, NV-5138 increased the phosphorylation of the mTORC1 downstream effector S6 (p-S6) in hippocampal and cortical lysates 2 hours post-administration (Figures 2A-D), indicating mTORC1 activation. Notably, NV-5138 also significantly decreased p-eIF2α (hippocampus, Figures 2A and 2B; cortex, Figurges 2C and 2D), consistent with the notion of crosstalk between the mTORC1 and the ISR in the brain. We confirmed this effect using IHC in excitatory neurons (Figures S4A-D). Intriguingly, quantitative IHC analysis revealed that p-S6 in *Fmr1*^−/y^ mice is elevated exclusively in excitatory but not inhibitory neurons (excitatory neurons, Figures S4E and S4F; inhibitory neurons, Figures S4G and S4H). Based on these findings, we hypothesized that increased activation of mTORC1 in excitatory neurons in *Fmr1*^−/y^ mice engenders dephosphorylation of eIF2α. To test this hypothesis, we suppressed mTORC1 in *Fmr1*^−/y^ animals by systemic administration of the mTORC1 inhibitor, CCI-779 (7.5 mg/kg, i.p., daily over 3 days), a rapamycin analog that was shown to cross the blood-brain barrier.^33, 34^ Inhibition of the enhanced mTORC1 activity in *Fmr1*^−/y^ mice corrected reduced p-eIF2α in the brain of these animals (hippocampus, Figures 2E and 2F; cortex, Figures 2G and 2H). In addition to pharmacological inhibition of mTORC1, we genetically downregulated mTORC1 selectively in excitatory neurons of *Fmr1*^−/y^ mice by deleting Raptor, a defining component of mTORC1, under the *Camk2α* promoter. Ablation of one allele of Raptor in excitatory neurons of *Fmr1*^−/y^ mice (*Fmr1*^−/y^; *raptor*^wt/f^; *Camk2α*^Cre^) corrected both the increased p-S6 (hippocampus and cortex, Figures S5A and S5B) and reduced p-eIF2α (hippocampus and cortex, Figures S5C and S5D) in excitatory neurons. Altogether, these results indicate that in *Fmr1*^−/y^ mice, a selective increase in mTORC1 activity in excitatory neurons leads to a decrease in p-eIF2α in this cell type.

**Figure 2.**
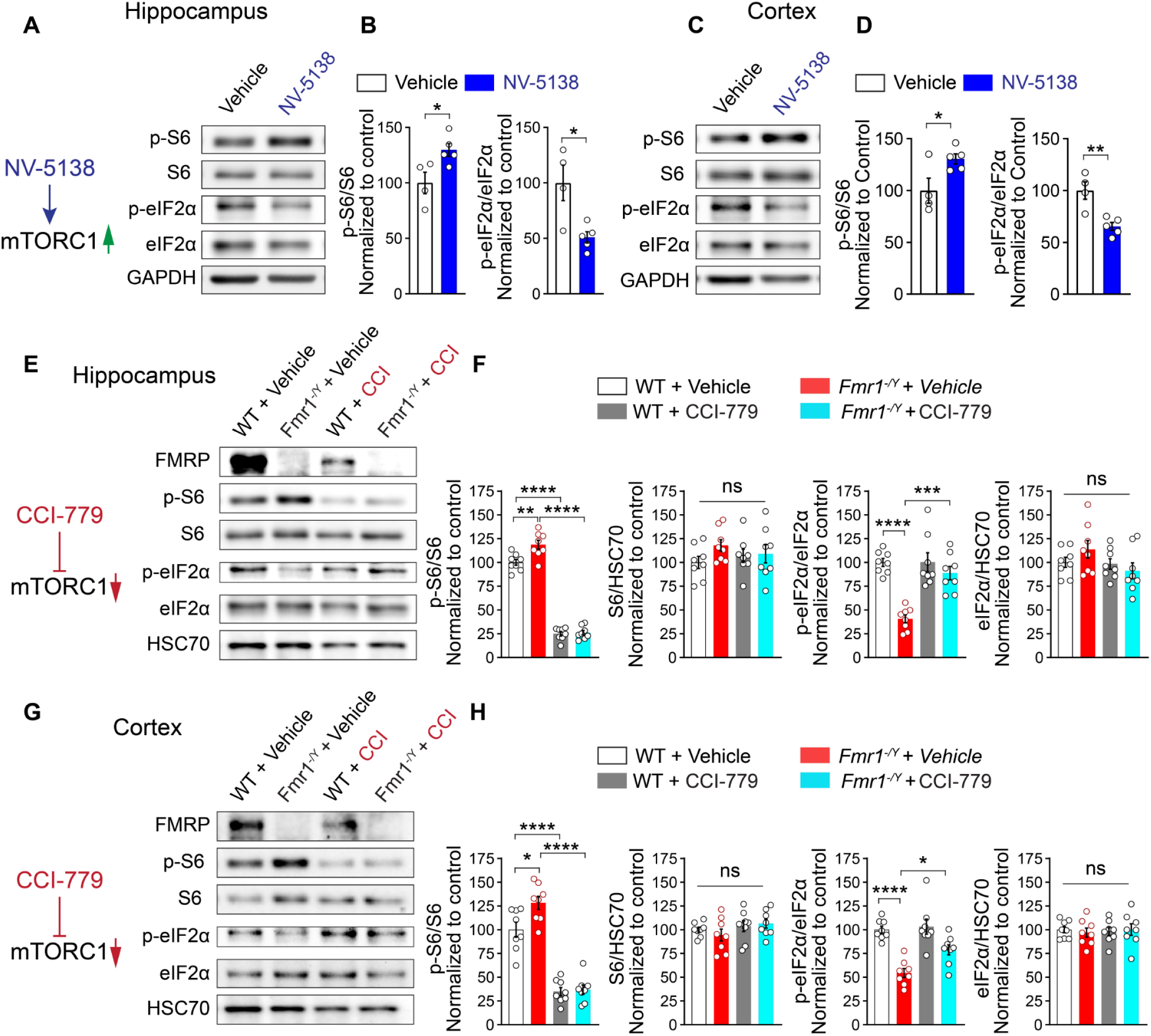
A crosstalk between mTORC1 and eIF2α pathways in the brain of *Fmr1^-/y^* mice. (A-D) Oral administration of NV-5138 results in elevated p-S6 and decrease in p-eIF2α in hippocampal and cortical lysates. (A and C) Brains were extracted from mice 2 hours after receiving vehicle (n = 4) or NV-5138 (160 mg/kg, n = 5)). NV-5138 causes an increase in p-S6/S6 and decrease in p-eIF2α in hippocampus (B, left, p-S6, *t* = 2.76, *p* = 0.0281; right, p-eIF2α, *t* = 3.211, *p* = 0.0148) and cortex (D, left, p-S6, *t* = 2.567, *p* = 0.0372; right, p-eIF2α, *t* = 4.039, *p* = 0.0049). Inhibition of mTORC1 with CCI-779 (7.5 mg/kg, daily over 3 days, i.p.) in the hippocampus (E) and cortex (G). (F) Quantifications of p-S6/S6 and p-eIF2α/eIF2α in hippocampus (F, p-S6/S6: *F*_3, 28_ = 228.4, *p* < 0.0001, WT + Vehicle versus *Fmr1*^-/y^ + Vehicle, *q*_28_ = 5.734, *p* = 0.0019; WT + Vehicle versus WT + CCI-779, *q*_28_ = 23.17, *p* < 0.0001; *Fmr1*^-/y^ + Vehicle versus *Fmr1*^-/y^ + CCI-779, *q*_28_ = 28.54, *p* < 0.0001, n = 8 per group; p-eIF2α/eIF2α, *F*_3, 28_ = 17.84, *p* < 0.0001, WT + Vehicle versus *Fmr1*^-/y^ + Vehicle, *q*_28_ = 8.838, *p* < 0.0001; *Fmr1*^-/y^ + vehicle versus *Fmr1*^-/y^ + CCI-779, *q*_28_ = 7.171, *p* = 0.0001). In cortex (H, p-S6/S6: *F*_3, 28_ = 57.32, *p* < 0.0001, WT + Vehicle versus *Fmr1*^-/y^ + Vehicle, *q*_28_ = 4.582, *p* = 0.0153; WT + Vehicle versus WT + CCI-779, *q*_28_ = 10.58, *p* < 0.0001; *Fmr1*^-/y^ + Vehicle versus *Fmr1*^-/y^ + CCI-779, *q*_28_ = 14.83, *p* < 0.0001; p-eIF2α/eIF2α: *F*_3, 28_ = 16, *p* < 0.0001, WT + Vehicle versus *Fmr1*^-/y^ + Vehicle, *q*_28_ = 8.082, *p* < 0.0001; *Fmr1*^-/y^ + vehicle versus *Fmr1*^-/y^ + CCI-779, *q*_28_ = 4.337, *p* = 0.0001. For E-H, n = 8 per group, one-way ANOVA followed by Tukey’s multiple comparisons post hoc test). No differences in S6 (hippocampus: *F*_3, 28_ = 1.002, *p* > 0.05 and cortex: *F*_3, 28_ = 1.033, *p* > 0.05) and eIF2α (hippocampus: *F*_3, 28_ = 1.652, *p* > 0.05 and cortex: *F*_3, 28_ = 0.1165, *p* > 0.05) were observed (One-way ANOVA). Each data point represents an individual animal. Data are presented as mean ± s.e.m. \**p* < 0.05, \*\**p* < 0.01, \*\*\*\**p* < 0.0001, ns, not significant. See also Figures S4, S5, S6, and S7.

### Increased levels of CaMK2α in *Fmr1*^−/y^ mice engender elevated mTORC1 activity in excitatory neurons

We next investigated the mechanism underlying excitatory neuron-specific activation of mTORC1 in *Fmr1*^−/y^ mice. Previous work has shown that CaMK2α can phosphorylate Homer, causing a disruption of Homer-mGluR5 interaction and thereby inducing mGluR5 activation.^35^ CaMK2α is an FMRP target,^36^ and its protein expression is increased in *Fmr1*^−/y^ mice,^35, 37^ leading to the hyperactivation of mGluR5. Elevated activity of mGluR5 in *Fmr1*^−/y^ mice signals to and enhances the activity of the PI3K-mTORC1 axis. Since CaMK2α is selectively expressed in excitatory but not inhibitory neurons, we hypothesised that increased expression of CaMK2α causes hyperactivation of mTORC1 selectively in excitatory neurons of *Fmr1*^−/y^ mice. Consistent with this hypothesis, ablation of one allele of *Camk2α* in *Fmr1*^−/y^ mice (*Fmr1*^-/y^; *Camk2*α^+/-^) corrected both the increased p-S6 and decreased p-eIF2α in *Fmr1*^−/y^ excitatory neurons (Figures S6A-D). To study whether the effect of FMRP ablation on p-S6 and p-eIF2α in excitatory neurons is cell type-autonomous, we deleted *Fmr1* selectively in excitatory neurons (*Fmr1*^f/y^; *EMX1*^Cre^). This manipulation was sufficient to cause an increase in p-S6 and decrease in p-eIF2α in excitatory neurons (hippocampus and cortex, Figures S7A-D). Conversely, rescue of *Fmr1* expression in excitatory neurons on *Fmr1*^−/y^ background^38^ (*Fmr1* conditional ON (cON); *EMX1*^Cre^) corrected the elevated p-S6 and normalized decreased p-eIF2α in this cell type (hippocampus and cortex, Figure S7E-H). Altogether, these experiments suggest that loss of FMRP causes mTORC1 hyperactivation and reduction in p-eIF2α via an excitatory neuron-dependent mechanism, mediated by increased CaMK2α protein expression.

### Decreased p-eIF2α in excitatory neurons causes phenotypes reminiscent of ASD

Elevated general protein synthesis is associated with ASD.^39–41^ To study whether the decrease in p-eIF2α, which causes an increase in general protein synthesis, is sufficient to engender autism-related phenotypes, we generated conditional knock-in (cKI) mice with reduced (∼50%) eIF2α phosphorylation selectively in excitatory neurons, to mimic the reduction of p-eIF2α in *Fmr1*^−/y^ mice. We crossed a transgenic mouse harboring one non-phosphorylatable Ser51Ala mutant *Eif2α* allele (*Eif2α*^S/A^) and the wild-type (WT) *Eif2α* transgene flanked by two *lox*-P sites^42^ with a mouse expressing Cre recombinase under the excitatory neuron-specific *Camk2α* promoter (referred to as *Eif2α*^S/A^ cKI^Camk2α^, Figure S8A). Cre recombinase in *Eif2α*^S/A^ cKI^Camk2α^ mouse leads to the excision of the WT transgene in excitatory neurons and induces eGFP expression. As expected, *Eif2α*^S/A^ cKI^Camk2α^ animals showed a reduction in p-eIF2α and an upregulation of protein synthesis (AHA-incorporation) in excitatory (p-eIF2α, Figure 3A; AHA, Figure 3B) but not inhibitory neurons (Figures S8B and S8C; analysis of total lysates in Figures 3C and 3D). Behavioral analysis showed that *Eif2α*^S/A^ cKI^Camk2α^ mice exhibit impaired social interaction in the novelty phase of the three-chamber social interaction test (Figures 3E and 3F) and reduced direct social interaction in the reciprocal social interaction test (Figure 3G). *Eif2α*^S/A^ cKI^Camk2α^ mice showed no deficits in novel object recognition (Figure S8D), olfactory discrimination (Figure S8E), and habituation/dishabituation (Figure S8F) tests. Moreover, *Eif2α*^S/A^ cKI^Camk2α^ mice exhibited increased self-grooming (Figure 3H) and repetitive marble burying behaviors (Figure 3I). No anxiety phenotype was found in the elevated plus maze (EPM, Figures S8G-I) in these mice. These findings demonstrate that the selective reduction of p-eIF2α in excitatory neurons is sufficient to engender altered social and repetitive/stereotypic behaviors, which are hallmarks of autism.

**Figure 3.**
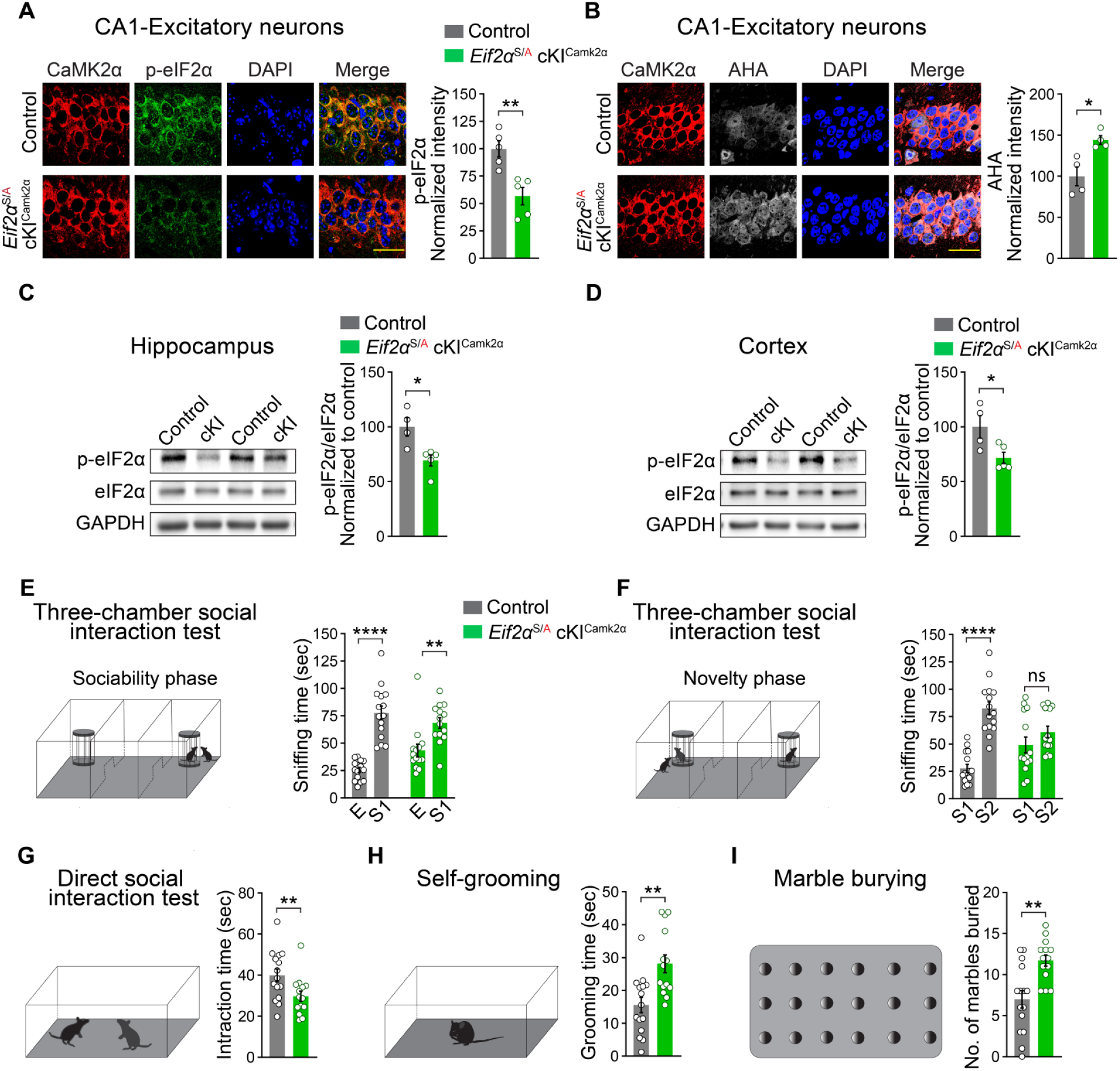
Mice with reduced phosphorylation of eIF2α in excitatory neurons exhibit autistic-like behaviors. **(A)** Immunostaining of hippocampal sections from *Eif2α*^S/A^ cKI^Camk2α^ mice shows a reduction in p-eIF2α in excitatory neurons (CaMK2α-positive, red) in CA1 area (control (n = 5) versus *Eif2α*^S/A^ cKI^Camk2α^ (n = 5), *t* = 4.038, *p* = 0.0037). (B) Global protein synthesis, measured by AHA incorporation, is increased in CA1 excitatory neurons in *Eif2α*^S/A^ cKI^Camk2α^ mice compared with control mice (control (n = 4) versus *Eif2α*^S/A^ cKI^Camk2α^ (n = 4), *t* = 3.504, *p* = 0.0128). (C, D) Immunoblotting (left) and quantification (right) show reduced p-eIF2α in *Eif2α*^S/A^ cKI^CamK2α^ (n = 5) compared with control (n = 4) mice in hippocampus (C, control versus *Eif2α*^S/A^ cKI^CamK2α^, *t* = 2.658, *p* = 0.0326), and cortex (D, control versus *Eif2α*^S/A^ cKI^CamK2α^, *t* = 3.296, *p* = 0.0132). *Eif2α*^S/S^ *Camk2*^Cre^ mouse line was used as control. (E and F) Three-chamber social interaction test in *Eif2α*^S/A^ cKI^Camk2α^ mice. (H) In the first 10-minutes phase of the test (sociability phase), both *Eif2α*^S/A^ cKI^Camk2α^ and control mice preferred cage containing stranger (S1) mouse over empty cage (E) (Chamber time effect: *F*_(1, 54)_ = 59.86, *p* < 0.0001; Controls, n = 15, E sniffing time versus S1 time, *t*_54_ = 7.528, *p* < 0.0001), Eif*2α*^S/A^ cKI^Camk2α^ mice (n = 14, E time versus S1 time, *t*_54_ = 3.485, *p* = 0.0020). Statistics are based on two-way ANOVA followed by Bonferroni’s post hoc test. In the second phase (F, novelty seeking phase), *Eif2α*^S/A^ cKI^Camk2α^ mice show no preference for novel mouse (S2) over familiar mouse (S1) (Chamber time effect; *F*_(1, 54)_ = 34.90, *p* < 0.0001; S1 time versus S2 time, *t*_54_ = 1.446, *p* > 0.05), contrary to control animals (S1 time versus S2 time, t_54_ = 7.006, *p* < 0.0001). Statistics are based on two-way ANOVA followed by Bonferroni’s post hoc test. Direct social interaction test (G) reveals that *Eif2α*^S/A^ cKI^Camk2α^ mice interact less with the stranger mouse than control mice (*t* = 2.49, *p* = 0.019). *Eif2α*^S/A^ cKI^Camk2α^ mice groom significantly more than controls (H, control (n = 15) versus *Eif2α*^S/A^ cKI^Camk2α^ (n = 14), *t* = 3.515, *p* = 0.0016) and bury more marbles in marble burying test (I, control (n = 15) versus *Eif2α*^S/A^ cKI^Camk2α^ (n = 14), *t* = 3.657, *p* = 0.0011). Student’s t-test was used in all panels except E and F. All data are shown as mean ± s.e.m. \**p* < 0.05, \*\**p* < 0.01, \*\*\**p* < 0.001, \*\*\*\**p* < 0.0001, and ns, not significant. Scale bars, 25 µm. See also Figure S8.

Next, we tested mice with whole-body reduction of p-eIF2α (∼50% reduction, *Eif2α*^S/A^ KI, Figure 4A, see Figures S9A-D for IHC analysis showing the reduction of p-eIF2α in excitatory and inhibitory neurons).^43^ Surprisingly, *Eif2α*^S/A^ KI animals showed no significant change in social interaction (three-chamber social interaction, Figures 4B and 4C; direct social interaction, Figure 4D), grooming (Figure 4E), marble burying (Figure 4F), or anxiety (EPM, Figures 4G-I). Notably, whereas mice with selective decrease in p-eIF2α in excitatory neurons (*Eif2α*^S/A^ cKI^Camk2α^) exhibited increased excitatory synaptic activity (mEPSC frequency) in pyramidal neurons (Figures S9E and S9F, also see ^44^), mice with whole-body reduction in p-eIF2α (*Eif2α*^S/A^ KI) showed no change in either synaptic excitation or inhibition (Figures S9G and S9H).

**Figure 4.**
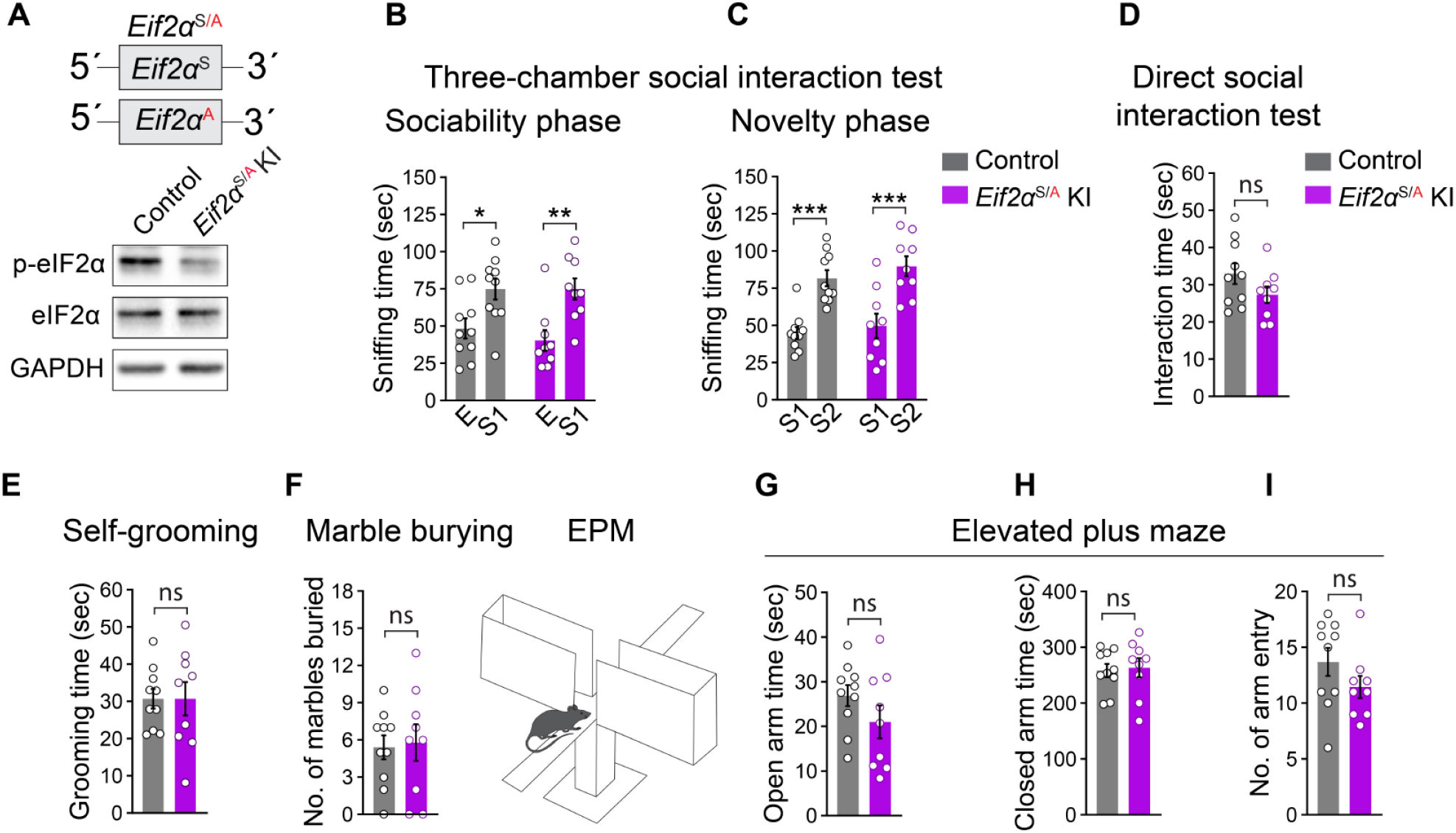
Heterozygous ablation of p-eIF2α in all cell types does not cause autism-like behaviors. (A, top) Schematic illustration of alleles in *Eif2α*^S/A^ knock-in (KI) mouse line. (A, bottom) Immunoblot shows a reduction in p-eIF2α in the brain of *Eif2α*^S/A^ KI mice. (B, C) three-chamber social interaction test. *Eif2α*^S/A^ KI mice show no deficits in sociability (B, *F*_(1, 34)_ = 0.337, *p* > 0.05; KI, n = 9; control, n = 10, sniffing time for empty cage (E) over stranger 1 (S1) for control, *t*_34_ = 2.766, *p* = 0.0182; KI mice *t*_34_ = 3.425, *p* = 0.0032, two-way ANOVA followed by Bonferroni’s multiple comparisons test) or novelty phase (C, *F*_(1, 34)_ = 0.082, *p* > 0.05, sniffing time for familiar mouse (S1) over novel mouse (S2) for control, *t*_34_ = 4.339, *p* = 0.0002; KI, *t*_34_ = 4.512, *p* = 0.0001, two-way ANOVA followed by Bonferroni’s multiple comparisons test). Similar to control animals (n = 10), *Eif2α*^S/A^ KI mice (n = 9) showed no impairments in direct social interaction (D, *t* = 1.561, *p* > 0.05, Student’s t-test), self-grooming (E, *t* = 0.005, *p* > 0.05, Student’s t-test), and marble burying (F, *t* = 0.218, p > 0.05, Student’s t-test). (G-I) No anxiety-like behavior was found in *Eif2α*^S/A^KI mice (open arm time, *t* = 1.373, *p* > 0.05; closed arm time, *t* = 0.2445, *p* > 0.05, and total number of arm entries, *t* = 1.396, *p* > 0.05, Student’s t-test). Each data point represents an individual animal. All data are presented as mean ± s.e.m. \**p* < 0.05, \*\**p* < 0.01, and \*\*\**p* < 0.001, and ns, not significant. See also Figure S9.

Taken together, the results demonstrate that the activity of the ISR is reduced selectively in excitatory neurons in *Fmr1*^−/y^ mice and is accompanied by an increase in general protein synthesis. Importantly, lowering p-eIF2α in excitatory neurons but not all cell types, engenders a synaptic excitation and inhibition imbalance and causes behaviors reminiscent of ASD.

### Rescue of *Fmr1*^−/y^ phenotypes by increasing p-eIF2α

To study whether normalization of the reduced p-eIF2α selectively in excitatory neurons in *Fmr1*^−/y^ mice could rescue increased protein synthesis and ASD-like features, we developed a genetic strategy to upregulate p-eIF2α only in excitatory neurons of *Fmr1*^−/y^ mice. To chronically increase p-eIF2α, we targeted the constitutively expressed regulatory protein CReP (encoded by the Protein Phosphatase 1 Regulatory Subunit 15B, *Ppp1r15b*), which forms an eIF2α phosphatase complex with PP1. GADD34, which can also recruit PP1 to eIF2α, is induced by the ISR and is therefore less suited for chronic modulation of p-eIF2α. We knocked down CReP in excitatory neurons using intracerebroventricular (i.c.v.) administration of adeno-associated virus (AAV) expressing a microRNA-adapted short hairpin RNA (shRNAmir) against *Ppp1r15b* under the excitatory neuron-specific *Camk2α* promoter (AAV9-Camk2α-GFP-*Ppp1r15b*-shRNAmir, Figures 5A and 5B). We used a viral titer that elevated p-eIF2α in excitatory neurons of *Fmr1*^−/y^ mice to the WT level (Figure 5C; characterization of the AAVs in Figures S10A-F). Notably, normalization of p-eIF2α in excitatory neurons of *Fmr1*^−/y^ mice corrected elevated protein synthesis (hippocampus, Figure 5D; cortex, Figure 5E).

**Figure 5.**
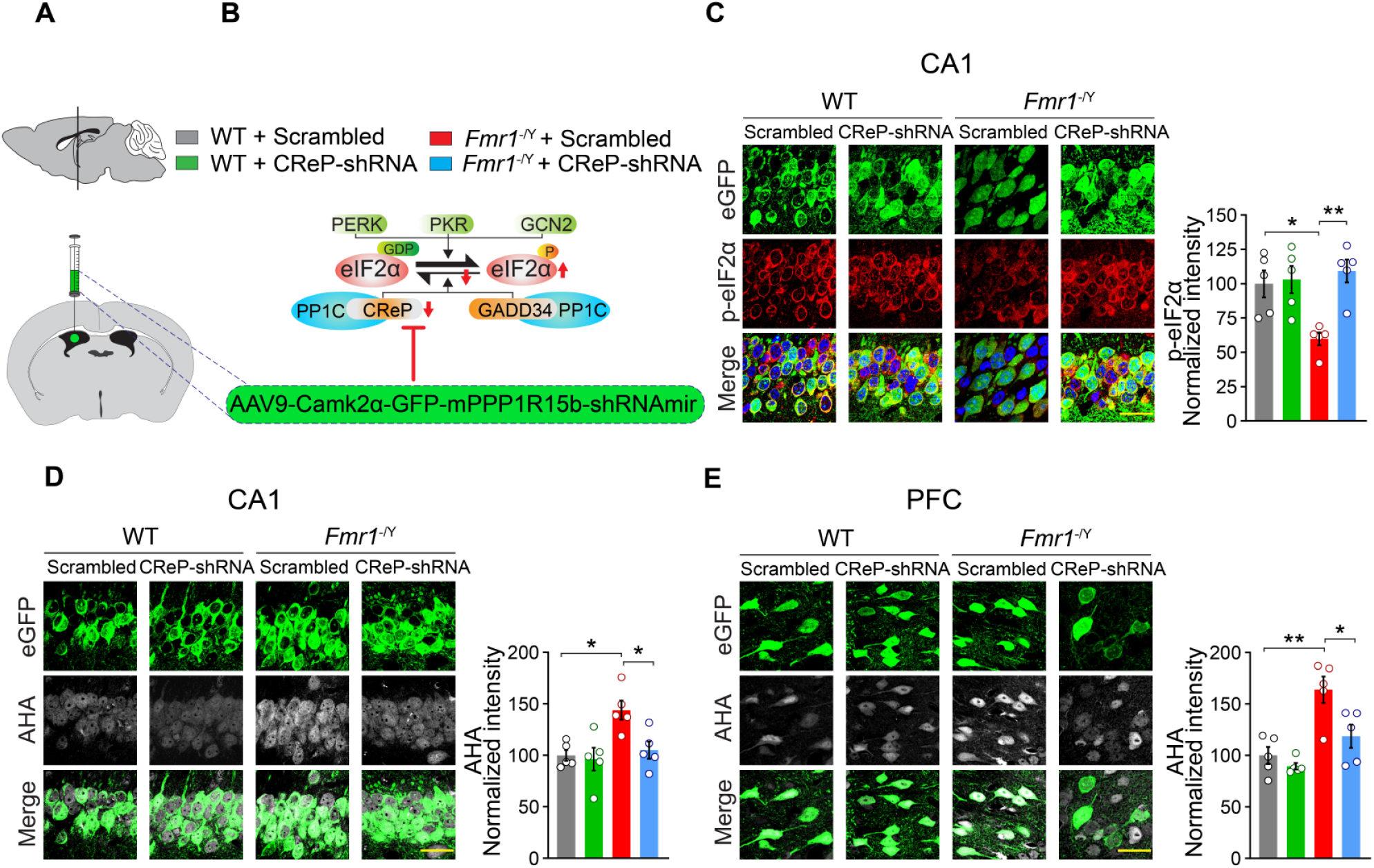
CReP-shRNA normalizes reduced eIF2α phosphorylation and corrects the elevated protein synthesis in excitatory neurons in *Fmr1*^-/y^ mice. (A) AAV9-Camk2α-GFP-*Ppp1r15b*-shRNAmir (CReP-shRNA) or scrambled AAV were delivered via intracerebroventricular (i.c.v.) injection to ablate CReP in excitatory neurons. AAVs were injected at postnatal day (PND) 28, and experiments were performed at PND P56. (B) In the brain, eIF2α is phosphorylated by PERK, PKR, and GCN2 kinases and dephosphorylated by PP1, which forms a complex with CReP or GADD34. (C) Immunostaining of hippocampal sections for p-eIF2α in wild-type and *Fmr1*^-/y^ animals injected with CReP-shRNA or scrambled AAVs. (D) CReP-shRNAmir increases the p-eIF2α in excitatory neurons (eGFP-positive) in CA1 of *Fmr1*^-/y^ mice to the WT level (*F*_3, 16_ = 7.054, *p* = 0.0031, *Fmr1*^-/y^ + CReP-shRNA versus *Fmr1*^-/y^ mice + Scrambled, *t*_16_ = 4.135, *p* = 0.0047; WT + Scrambled versus *Fmr1*^-/y^ + Scrambled, *t*_16_ = 3.352, *p* = 0.0243, n = 5 for all groups. Statistics are based on one-way ANOVA followed by Tukey post hoc comparisons. AHA incorporation (grey) in excitatory neurons (eGFP-positive, green) in CA1 (D) and PFC (E). CReP-shRNA AAV normalizes the elevated protein synthesis in excitatory neurons in *Fmr1*^-/y^ mice in CA1 (D, *F*_3, 16_ = 6.286, *p* = 0.005, WT + Scramble versus WT + CReP-shRNA, *q*_16_ = 0.411, *p* > 0.05; WT + Scrambled versus *Fmr1*^-/y^ + Scrambled, *q*_16_ = 5.004, *p* = 0.0131; *Fmr1*^-/y^ + Scrambled versus *Fmr1*^-/y^ + CReP-shRNA, *q*_16_ = 4.424, *p* = 0.0297, one-way ANOVA followed by Tukey post hoc comparisons, n = 5 for each group), and PFC (E, *F*_3, 16_ = 11.62, *p* = 0.0003, WT + Scrambled versus WT + CReP-shRNA, *q*_16_ = 1.085, p > 0.05; WT + Scrambled versus *Fmr1*^-/y^ + Scrambled, *q*_16_ = 6.627, *p* = 0.0013; *Fmr1*^-/y^ + Scrambled versus *Fmr1*^-/y^ + CReP-shRNA, *q*_16_ = 4.688, *p* = 0.0205, one-way ANOVA followed by Tukey’s multiple comparisons test, n = 5 for each group). Each data point represents individual animal. Scale bar, 25 μm. All data are shown as mean ± s.e.m. \**p* < 0.05, \*\**p* < 0.01, ns, not significant. See also Figure S10.

We next studied whether correction of p-eIF2α in excitatory neurons rescues core *Fmr1*^−/y^ mice phenotypes. We first examined the effect of normalizing p-eIF2α in excitatory neurons on the exaggerated group 1 metabotropic glutamate receptor (mGluR)-dependent long-term depression (LTD), a key feature of *Fmr1*^−/y^ mice.^45^ Whereas *Fmr1*^−/y^ mice injected with the control AAV (expressing scrambled sequence) exhibited enhanced LTD in response to DHPG administration, *Fmr1*^−/y^ mice injected with AAV-CReP-shRNAmir exhibited intact mGluR-LTD (Figures 6A-D), demonstrating correction of this phenotype.

**Figure 6.**
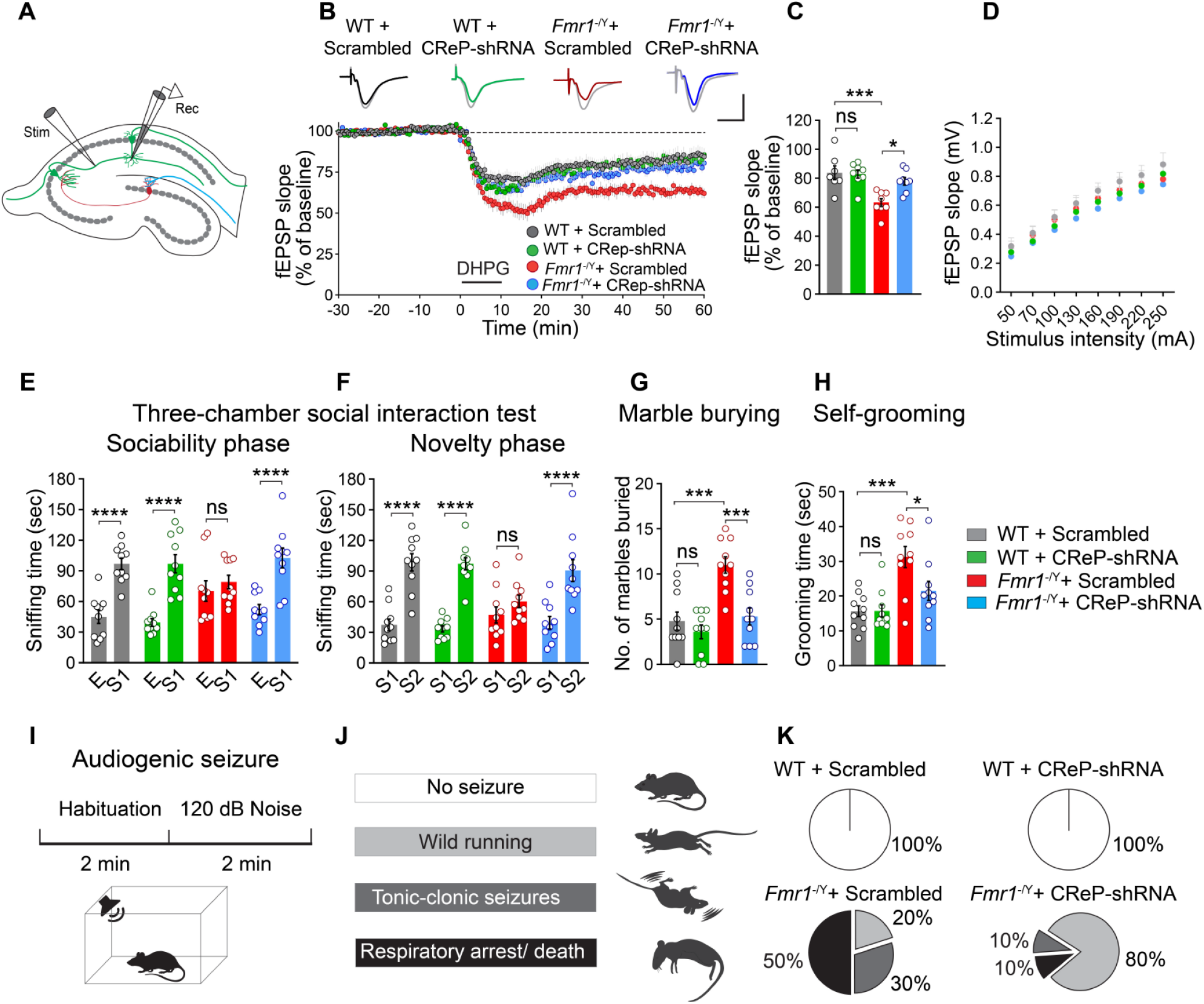
Correction of eIF2α phosphorylation in excitatory neurons rescues pathological phenotypes in *Fmr1*^-/y^ mice. (A-D) Ablation of CReP partially rescues exaggerated mGluR-LTD. (A) CReP-shRNA or Scrambled AAV was injected i.c.v. at postnatal day (PND) 1-2 and recording was performed at PND 28-35. (B) Representative traces from 4 groups (baseline in grey and recording at 1 hour in solid colors). (Bottom) LTD was induced by application of DHPG (50 µM for 10 min). Field excitatory postsynaptic potentials (fEPSP) were recorded over a 60-min period after DHPG-induced LTD. (C) The quantification of the fEPSP slope (% of baseline) during the last 10 min of the recording. CReP-shRNA AAVs reduced the exaggerated LTD in *Fmr1*^-/y^ mice (*F*_3, 27_ = 8.587, *p* = 0.0004, WT + Scrambled (n = 7 slices from 7 mice) versus *Fmr1*^-/y^ + Scrambled (n = 8 slices from 8 mice), *q*_27_ = 6.196, *p* = 0.0009; *Fmr1*^-/y^ + Scrambled versus *Fmr1*^-/y^ + CReP-shRNA (n = 8 slices from 8 mice), *q*_27_ = 4.580, *p* = 0.0158; one-way ANOVA followed by Tukey’s multiple comparisons post hoc test). (D) No differences in input/output responses (*F*_21, 182_ = 0.643, p > 0.05, one-way ANOVA). (E, F) Three-chamber social interaction test, n = 10 per group. (E) In sociability phase, CReP-shRNA AAV rescued the time *Fmr1*^-/y^ mice interact with stranger animal (S1) over empty cage (E) (Chamber time effect, *F*_(1, 72)_ = 66.10, *p* < 0.0001; *Fmr1*^-/y^ + CReP-shRNA, *t*_72_ = 4.859, *p* < 0.0001; WT + Scrambled, *t*_72_ = 5.005, *p* < 0.0001; *Fmr1*^-/y^ + Scrambled, *t*_72_ = 0.849, *p* > 0.5; WT + CReP-shRNA, *t*_72_ = 5.546, *p* < 0.0001, one-way ANOVA followed by Bonferroni’s multiple comparisons test). (F) In the novelty seeking phase, *Fmr1*^-/y^ + CReP-shRNA group spent significantly more time interacting with novel moues (S2) than familiar one (S1) (Chamber time effect, *F*_(1, 72)_ = 90.46, *p* < 0.0001; Fmr*1*^-/y^ + CReP-shRNA*, t*_72_ = 5.137, *p* < 0.0001; WT + Scrambled, *t*_72_ = 6.147, *p* < 0.0001; *Fmr1*^-/y^ + Scrambled, *t*_72_ = 1.343, *p* > 0.5; WT + CReP-shRNA, *t*_72_ = 6.394, *p* < 0.0001, one-way ANOVA followed by Bonferroni’s multiple comparisons test). (G) In marble burying test (n = 10 per group), CReP-shRNA AAV reduced the number of buried marbles in *Fmr1*^-/y^ mice (*F*_3, 36_ = 13.20, *p* < 0.0001, WT + Scrambled versus *Fmr1*^-/y^ + Scrambled, *q*_36_ = 6.836, *p* = 0.0001; *Fmr1*^-/y^ + Scrambled vs. *Fmr1*^-/y^ + CReP-shRNA, *q*_36_ = 6.285, *p* < 0.0005, One-way ANOVA followed by Tukey’s multiple comparisons test). (H) CReP-shRNA AAV reduced the time *Fmr1*^-/y^ mice spent grooming (*F*_3, 36_ = 9.602, *p < 0.0001,* WT + Scrambled versus *Fmr1*^-/y^ + Scrambled, *q*_36_ = 6.587, *p* = 0.0002; *Fmr1*^-/y^ + Scrambled versus *Fmr1*^-/y^ + CReP-shRNA, *q*_36_ = 4.108, *p* = 0.304, One-way ANOVA, followed by Tukey’s multiple comparisons test). For E-H, AAVs were injected i.c.v. at postnatal day 28, and experiments were performed starting PND P56. (I-K) CReP-shRNA AAV alleviated audiogenic seizures (AGS) in *Fmr1*^-/y^ mice. (I) AGS induction protocol (top), and the apparatus (bottom) composed of a soundproof box and a speaker to generate 120 dB noise. (J) Different levels of seizure upon exposure to 120 dB noise ranging from no seizure to wild running (WR), tonic-clonic (TC), and severe seizure that leads to respiratory arrest and death (RA). (K) Unlike WT + Scrambled (n = 10) and WT + CReP-shRNA (n = 10) groups, which showed no seizure, 20% of *Fmr1*^-/y^ mice (n = 10) experience wild running, 30% tonic-clonic and 50% of them showed the severe form of the seizure, respiratory arrest, and death. CReP-shRNA AAV significantly reduced the severity of the seizure in *Fmr1*^-/y^ + CReP-shRNA (n = 10), (*X*^2^ (9)=545.3, *p* < 0.0001, Fisher’s exact test). For I-K, AAVs were injected i.c.v. at postnatal day 1-2, and experiments were performed at PND P28-35. Each data point represents individual animal. All data are shown as mean ± s.e.m. \**p* < 0.05, \*\*\**p* < 0.001, \*\*\*\**p* < 0.0001, ns, not significant. See also Figure 11.

We next investigated whether correction of p-eIF2α rescues altered social and repetitive/stereotypic behaviors in *Fmr1*^−/y^ mice. Remarkably, normalization of p-eIF2α in excitatory neurons of *Fmr1*^−/y^ mice rescued social deficits in the three-chamber social interaction test (Figures 6E and 6F) and reversed the exaggerated marble burying (Figure 6G) and grooming (Figure 6H) behaviors. Moreover, the audiogenic seizure phenotype of *Fmr1*^−/y^ mice was substantially alleviated upon correction of p-eIF2α (Figures 6I-K). Altogether, these results demonstrate that normalization of reduced p-eIF2α in excitatory neurons of *Fmr1*^−/y^ mice corrects impaired social interaction and repetitive behaviors and ameliorates the audiogenic seizure phenotype.

## Discussion

Dysregulation of protein synthesis is a paramount pathophysiological mechanism in FXS and several other neurodevelopmental disorders.^2, 4, 39^ Our study is the first to show that decreased activity of the ISR, associated with reduced p-eIF2α and elevated general protein synthesis, is sufficient to engender social interaction deficits and repetitive behaviors. Moreover, we demonstrate that this mechanism critically contributes to core ASD-like phenotypes in a mouse model of FXS. Correction of reduced p-eIF2α in excitatory neurons of *Fmr1*^−/y^ mice not only normalized enhanced protein synthesis, but also rescued social interaction deficits, repetitive behavior, and audiogenic seizure phenotypes.

Our study shows that p-eIF2*α* is decreased and protein synthesis is increased in excitatory but not inhibitory neurons in a mouse model of FXS. Selective ablation of p-eIF2*α* in excitatory neurons leads to their hyperactivation.^44^ This might cause an imbalance between excitatory and inhibitory neuronal circuit activity resulting in abnormal neuronal functioning and social deficits, which are commonly observed in ASD mouse models and individuals with ASD.^46–48^ This notion is supported by the finding that reduction of p-eIF2*α* in excitatory neurons (*Eif2α*^S/A^ cKI^Camk2α^) but not all cell types (*Eif2α*^S/A^ KI), engenders an imbalance between excitatory and inhibitory synaptic inputs and autistic features.

The ISR pathway is differentially modulated in distinct neuronal cell types to control physiological processes such as memory formation^44^ and skill learning.^49^ Our work reveals cell-type-specific alterations of the ISR in a neurodevelopmental disorder. Moreover, we show that reduced activity of the ISR in excitatory neurons is sufficient to cause phenotypes reminiscent of ASD. These findings highlight the need to assess translational control pathways and rates of protein synthesis in disease states in a cell-type-specific manner as their evaluation in total cell lysates might dilute the effect of cell-type-specific alterations and mask underlying mechanisms. These results also imply that cell-type-specific interventions, targeting cell populations in which the ISR is dysregulated, may offer more effective treatments compared with approaches targeting all cell types.

Our work shows that the activities of the two main translational control mechanisms, mediated via the mTORC1 and the ISR, are coordinated in the brain (Figure 2 and Figure S11). We found that stimulation of mTORC1 in the brain leads to a downregulation in p-eIF2α, and mTORC1 inhibition in *Fmr1*^−/y^ mice normalizes p-eIF2α, suggesting that enhanced mTORC1 activity contributes to reduced ISR in FXS. Future studies should investigate the molecular mechanism by which mTORC1 activation downregulates p-eIF2α in both neuronal and non-neuronal cells. Intriguingly, mTORC1 activity is increased in excitatory but not inhibitory neurons where it causes a selective downregulation of p-eIF2α. The neuronal cell type-specific interplay between mTORC1 and ISR underscores the intricacy of translational control mechanisms in the brain, demonstrating how changes in one signaling pathway can contribute to phenotypes by altering the homeostasis of another signaling cascade.

Ablation of *Fmr1* in excitatory neurons is sufficient to cause core FXS phenotypes such as audiogenic seizures,^50, 51^ and social interaction deficits,^52^ indicating a preeminent role for excitatory neurons in FXS. Yet, changes in inhibition in *Fmr1*^−/y^ mice and their important roles in FXS pathophysiology are also well documented.^1, 53–55^ These changes are likely mediated via eIF2α-independent mechanisms.

In summary, we uncovered the role of excitatory neuron-specific suppression of the ISR pathway in engendering core ASD-like phenotypes in *Fmr1*^−/y^ mice.

## Acknowledgments

This work was supported by a grant from the Simons Foundation (SFARI, award #611773), the Canadian Institutes of Health Research (CIHR, PJT-162412), and the Azrieli Centre for Autism Research (ACAR) to A.K., and a Hellenic Foundation for Research and Innovation (H.F.R.I.) under the ‘2nd Call for H.F.R.I. Research Projects to support Faculty Members & Researchers’ (Project Number: 2556) grant to C.G.G. J.C.L. is supported by the Canada Research Chair in Cellular and Molecular Neurophysiology (CRC 950-231066). NIH Fragile X Center, U54HD104461 (K.H. and J.G.). M.H. was supported by the Brain Canada and Transforming Autism Care Consortium (TACC) graduate fellowships. We thank Gottfried Simbriger for helping with the uORF analysis. We thank Dr. David L. Nelson (Baylor College of Medicine, Houston) for kindly providing *Fmr1*^f/f^,^56^ and *Fmr1* conditional on (cON) mice.^38^

## Author contributions

M.H., N.S., C.G.G., and A.K. conceived the project, designed experiments, and supervised the research. M.H. performed immunohistochemistry and biochemistry analyses, behavioral experiments, and LTD recordings. C.T.P. and J.C.L. assisted with electrophysiological studies. M.L., M.A., E.S., C.W., P.M.R., E.N., I.G., K.C.L., and J.P. helped with data analysis. K.N. and M.C.M. assisted with behavioral experiments and data interpretation. J.R.G., K.H., R.J.K., V.S., and R.S. assisted with the generation of transgenic animals. K.S. assisted with the analysis of previous TRAP studies. R.J.H., T.D.B., and V.M-C., assisted with human brain tissue analysis. C.R., G.P., and T.M.D. helped with iPSC experiments. M.H., N.S., C.G.G., and A.K. wrote the manuscript.

---

**Figure S1.**
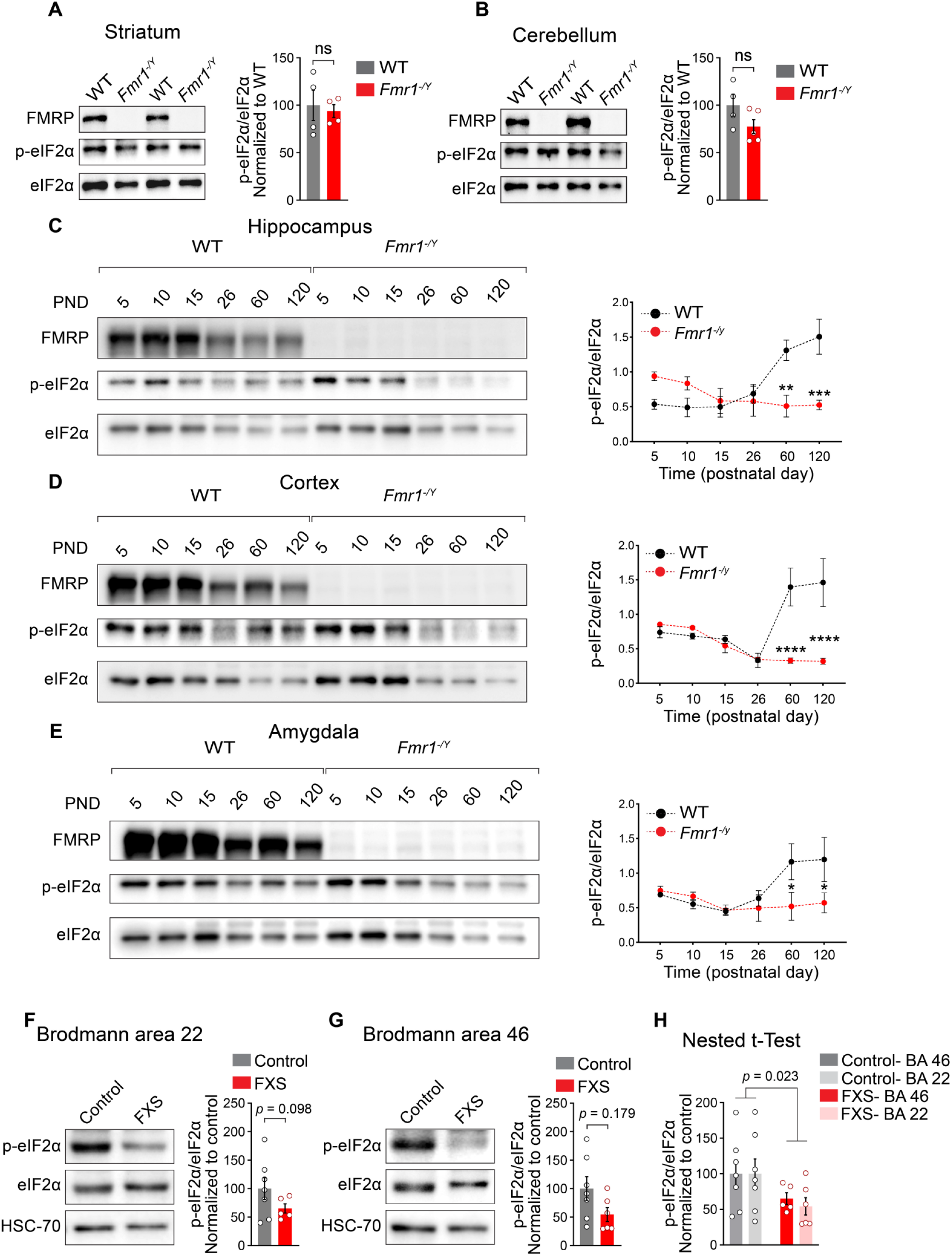
Expression of p-eIF2α in *Fmr1*^-/y^ mice during development and in post-mortem brain tissues collected from patients with fragile X syndrome. (A, B) Immunoblotting (left) and quantification (right) show no differences in p-eIF2α levels in *Fmr1*^-/y^ mice (n = 4-5) compared with WT mice (n = 4) in striatum (A, WT versus *Fmr1*^-/y^, *t* = 0.3457, *p* > 0.05) and cerebellum (B, WT versus *Fmr1*^-/y^, *t* = 1.697, *p* > 0.05). (C-E) Immunoblotting (left) and quantification (right) show a significant reduction in p-eIF2α levels in *Fmr1*^-/y^ mice at postnatal day (PND) 60 and 120 in lysates from hippocampus (C, *F*_5, 36_ = 7.687, *p* < 0.0001; WT (n = 4) versus *Fmr1*^-/y^ (n = 4); PND 60 (*t*_36_ = 3.792, *p* = 0.0033), and PND 120 (*t*_36_ = 4.64, *p* = 0.0003), cortex (D, *F*_5, 36_ = 9.093, p < 0.0001, WT (n = 4) versus *Fmr1*^-/y^ (n = 4); PND 60 (*t*_36_ = 5.403, *p* < 0.0001), and PND 120 (*t*_36_ = 5.776, *p* < 0.0001), and amygdala (E, *F*_5, 36_ = 2.387, p = 0.0572, WT (n = 4) versus *Fmr1*^-/y^ (n = 4); PND 60 (*t*_36_ = 2.886, *p* = 0.0394), and PND 120 (*t*_36_ = 2.809, *p* = 0.0479). For all groups, two-way ANOVA followed by Bonferroni’s multiple comparisons tests were performed. (F-H) Representative immunoblotting (left) and quantification (right) exhibit a trend toward a reduction in p-eIF2α levels in Brodmann areas (BA) 46 (F, *t* = 1.443, *p* = 0.1796) and 22 (G, *t* = 1.802, *p* = 0.0989) of the post-mortem brain tissues from fragile X syndrome patients (n = 5-6) compared with healthy controls (n = 7), Student’s t-test was performed for both areas. (H) Nested t-test analysis of combined BA 46 and BA 22 samples revealed a significant decrease in p-eIF2α in FXS patients compared with controls (*t* = 2.421, *p* = 0.0238, Nested t-test analysis). All data are shown as mean ± s.e.m. \**p* < 0.05, \*\**p* < 0.01, \*\*\**p* < 0.001, \*\*\*\**p* < 0.0001, ns, not significant.

**Figure S2.**
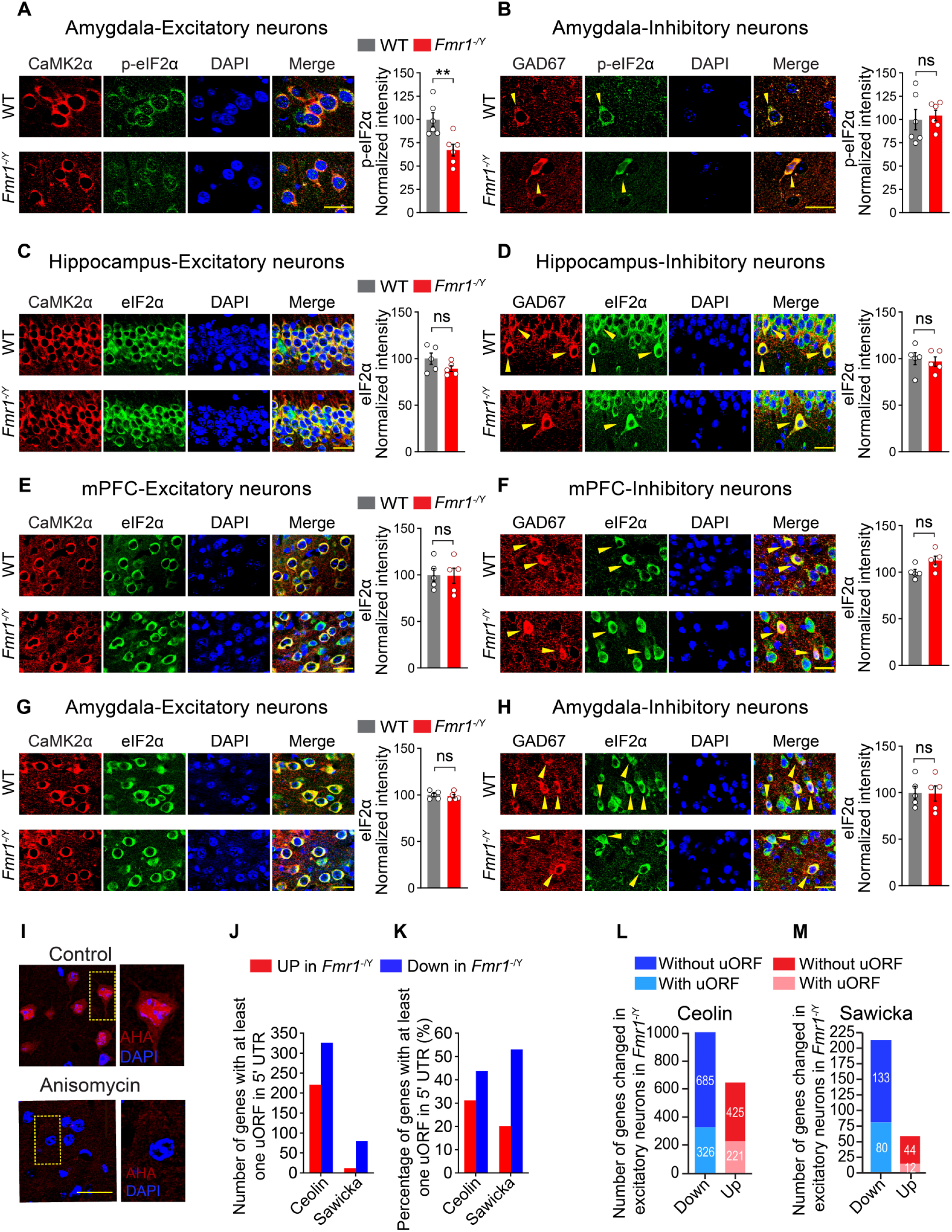
The expression of total eIF2α in different brain areas of *Fmr1*^-/y^ mice. (A) Immunostaining and quantification of p-eIF2α expression in amygdala excitatory neurons in *Fmr1^-/y^* (n = 6) compared with WT (n = 6) mice show decreased p-eIF2α (A, *t* = 3.409, *p* = 0.0067, Student’s t-test). No difference between *Fmr1^-/y^* (n = 6) and WT (n = 6) animals was observed in inhibitory neurons in amygdala (B, *t* = 0.345, *p* > 0.05, Student’s t-test). (C-H) Immunostaining and quantification of total eIF2α expression in excitatory and inhibitory neurons in CA1, PFC, and amygdala showing no change in eIF2α in excitatory neurons of *Fmr1^-/y^* mice (n = 5) compared with WT animals (n = 5) (C, CA1, *t* = 1.565*, p* > 0.05; E, PFC, *t* = 0.086*, p* > 0.05; G, amygdala, *t* = 0.534*, p* > 0.05, Student’s t-test) and in inhibitory neurons (D, CA1, *t* = 0.371, *p* > 0.05; E, PFC, *t* = 2.252, *p* > 0.05; F, amygdala, *t* = 0.3578, *p* > 0.05, Student’s t-test). Yellow arrows mark inhibitory neurons. (I) AHA incorporation into newly synthesized polypeptides is sensitive to anisomycin. (I, top, AHA in the control section is incorporated into newly synthesized proteins, and (bottom) pre-treatment with anisomycin blocks AHA incorporation. (J-M) Analysis of previously published TRAP data from excitatory neurons of *Fmr1^-/y^*mice (Ceolin ^11^, Sawicka ^12^) shows a reduction in genes harboring uORF (*p* < 0.0001 for both datasets, two-sided Fisher’s exact test). The bar graphs demonstrate the number (J) and the percentage (K) of genes containing uORF in their 5′ UTR, and (L, M) the total number of upregulated and downregulated genes with and without uORF in the 5′ UTR in excitatory neurons in *Fmr1^-/y^* mice. Each data point represents an individual animal. All data are presented as mean ± s.e.m. \**p* < 0.05, \*\**p* < 0.01, and ns, not significant. Scale bars, 25 µm.

**Figure S3.**
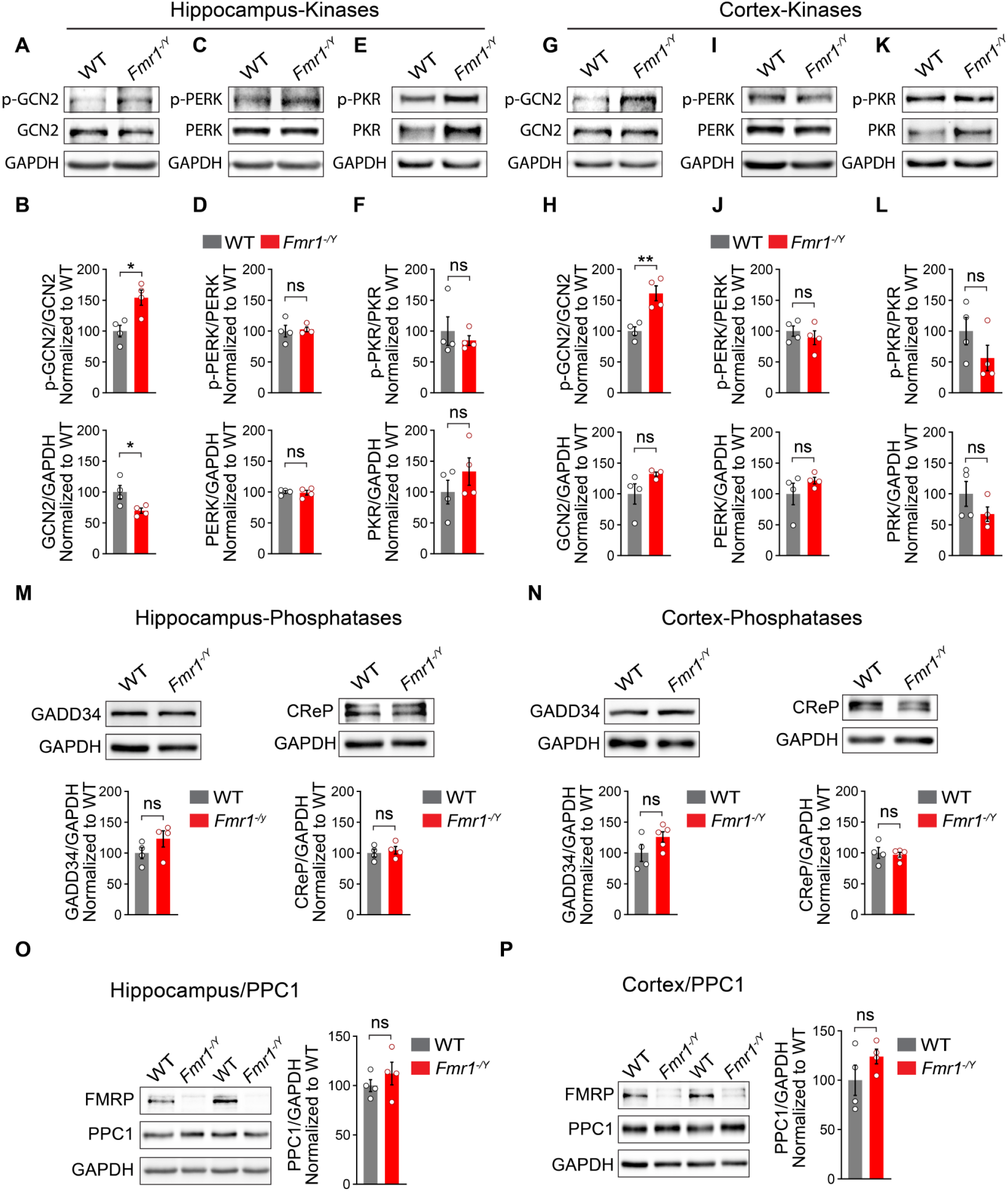
eIF2α phosphatases and kinases in the brain of *Fmr1^-/y^* mice. (A-L) Representative immunoblots and quantifications of p-GCN2, p-PERK, and p-PKR in hippocampal and cortical lysates extracted from *Fmr1^-/y^* (n = 4) and WT animals (n = 4) show increased p-GCN2 in the hippocampus (B, *t* = 3.469, *p* = 0.0133) and cortex (H, *t* = 4.417, *p* = 0.0045) of *Fmr1^-/y^* animals. No differences in p-PERK and p-PKR in the hippocampus (D, F, top, *p* > 0.05) and cortex (J, L, top, *p* > 0.05). No differences in total PERK and PKR in the hippocampus (D, F, bottom, *p* > 0.05) and cortex (J, L, bottom, *p* > 0.05). The ratio of total GCN2 to loading control in hippocampus lysates of *Fmr1^-/y^* was lower than in WT mice (B, bottom, *t* = 2.656, *p* = 0.0377). Representative images and quantification of GADD34 and CReP in hippocampal (M) and cortical (N) lysates. The levels of GADD34 (hippocampus: M, right, *p* > 0.05; cortex: N, right, *p* > 0.05) and CReP (hippocampus: M, left, *p* > 0.05; cortex: N, left, *p* > 0.05) in *Fmr1^-/y^* (n = 4) hippocampal and cortical lysates are not different comparing with WT (n = 4) animals. No differences were found in the level of catalytic subunit of protein phosphatase 1 (PP1C) in hippocampal (O, *p* > 0.05) or cortical (P, *p* > 0.05) lysates extracted from *Fmr1*^-/y^ mice versus WT animals. Student’s t-test was performed for all experiments. Each data point represents an individual animal. Data are presented as mean ± s.e.m. \**p* < 0.05, \*\**p* < 0.01, ns, not significant.

**Figure S4.**
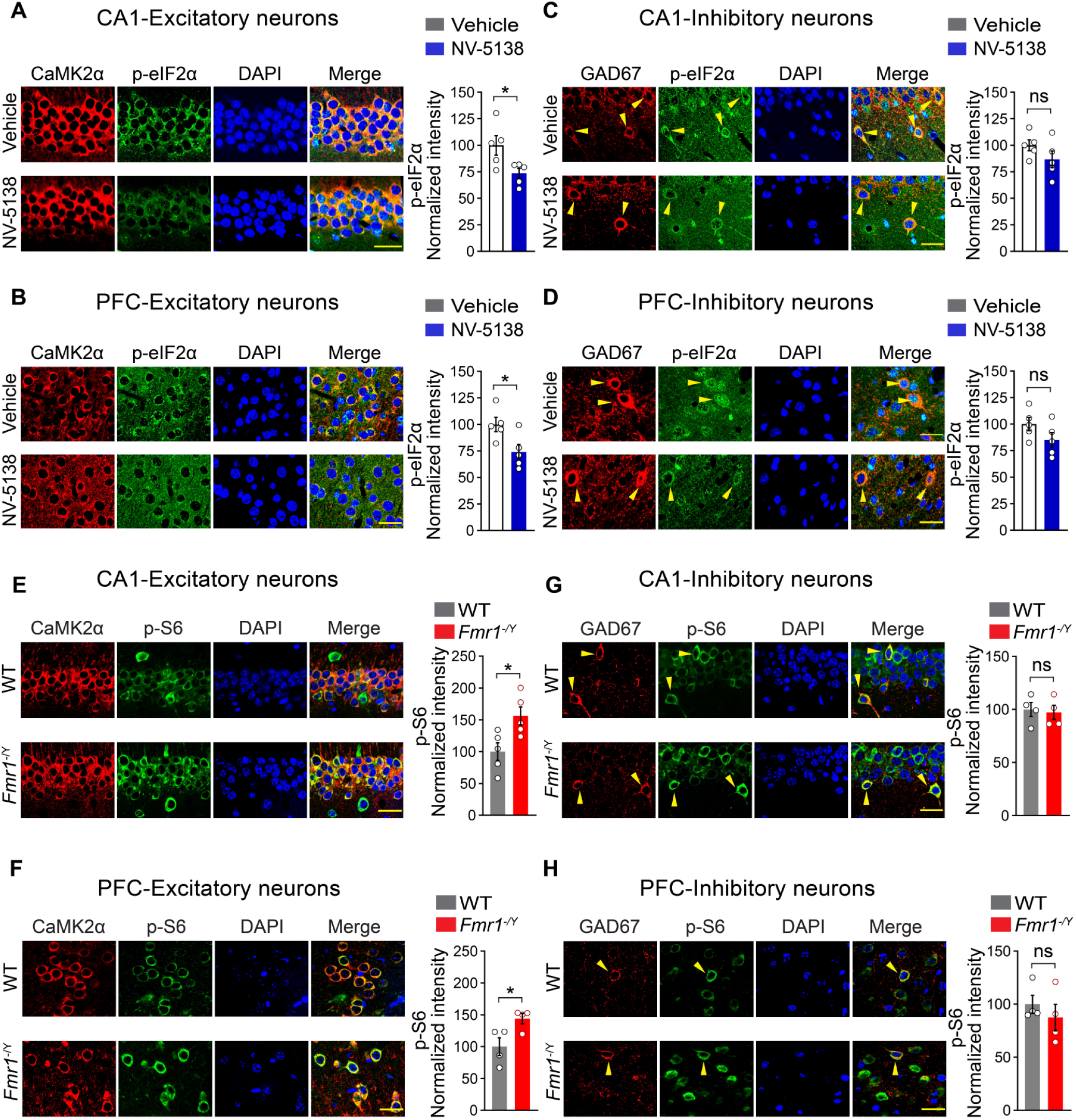
Elevated p-S6 in excitatory neurons in the brain of *Fmr1*^-/y^ mice. (A-D) Representative images and quantification of p-eIF2α expression in excitatory and inhibitory neurons in the hippocampus and PFC of mice show that NV-5138 treatment decreases p-eIF2α in excitatory neurons compared with vehicle (A, hippocampus, *t* = 2.568, *p* = 0.0332 ; B, PFC, *t* = 2.665, *p* = 0.0286). NV-5138 does not affect p-eIF2α in inhibitory neurons in the hippocampus and PFC (C, hippocampus, *p* > 0.05; D, PFC, *p* > 0.05), n = 5 for all groups. (E-H) Representative images and quantification of p-S6 in excitatory and inhibitory neurons from *Fmr1^-/y^* and WT mice show that p-S6 is higher in *Fmr1^-/y^* excitatory neurons in hippocampus (E, *t* = 2.845, *p* = 0.0217, n = 5 for both groups), and PFC (F, *t* = 2.714, *p* = 0.0349, n = 4 for either group). No differences were observed in inhibitory neurons in p-S6 levels in hippocampus (G, *t* = 0.293, *p* > 0.05, n = 4 for both groups) and PFC (H, *t* = 0.833, *p* > 0.05) versus WT mice (n = 4). Student’s t-test was performed for all analyses. Yellow arrows show inhibitory neurons. Each data point represents an individual animal. All data are presented as mean ± s.e.m. \**p* < 0.05, ns, not significant. Scale bars, 25 µm.

**Figure S5.**
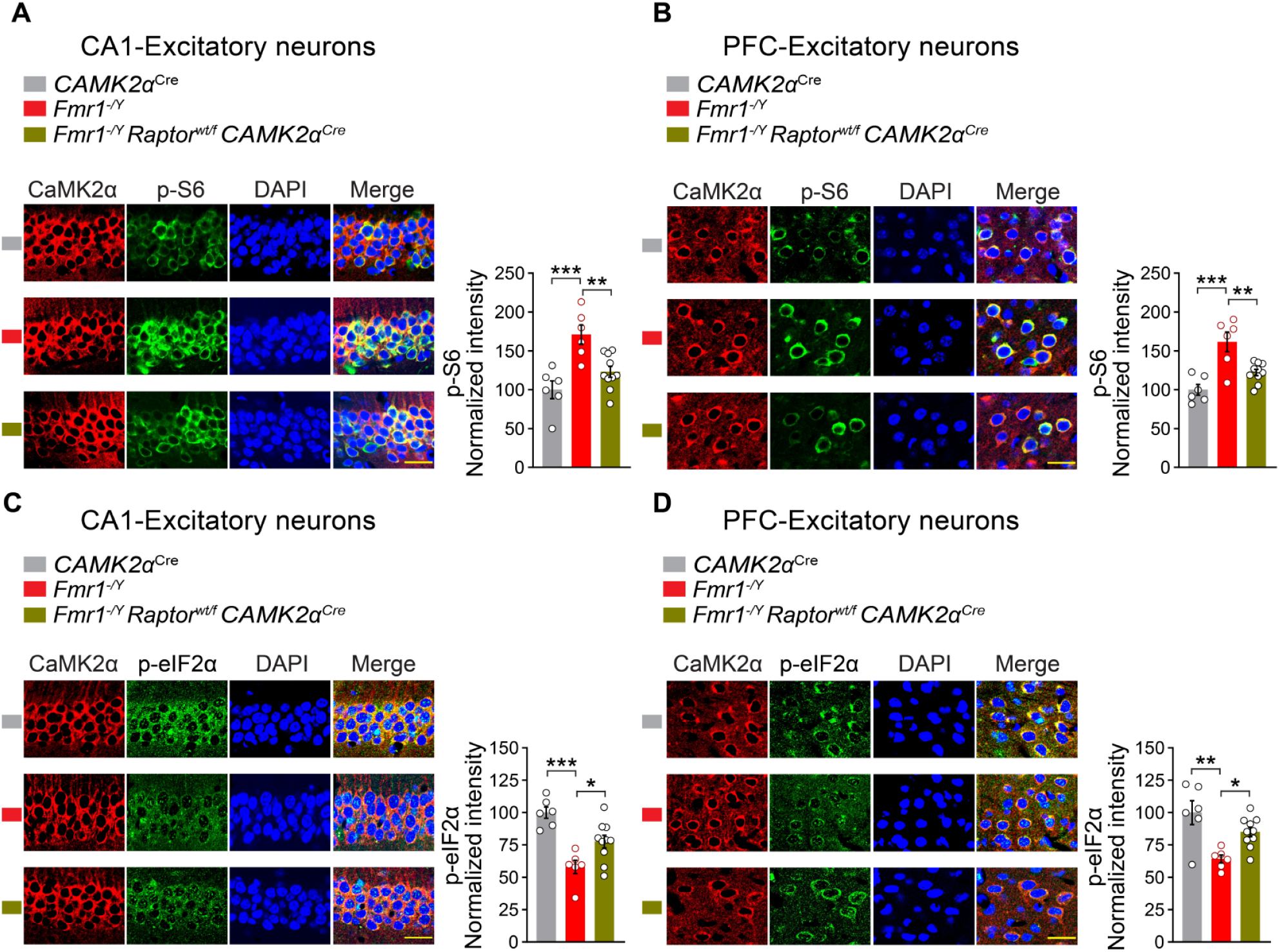
Deletion of one allele of Raptor in excitatory neurons rescues elevated p-S6 and reduced p-eIF2α in these neurons in *Fmr1*^-/y^ mice. (A-D) Representative images and quantification of p-S6 and p-eIF2α in excitatory neurons in the hippocampus and PFC of *Fmr1*^-/y^ mice with ablation of one allele of Raptor (*Fmr1*^-/y^ *Raptor*^wt*/*f^ *CaMK2α*^Cre^). Downregulation of Raptor rescues p-S6 (A, hippocampus; *F*_2, 19_ = 11.58, *p* = 0.0005, *CaMK2α*^Cre^ versus *Fmr1*^-/y^, *p* = 0.0005, *q*_19_ = 6.624; *CaMK2α*^Cre^ versus *Fmr1*^-/y^ *Raptor*^wt/f^ *CaMK2α*^Cre^, *p* > 0.05; *Fmr1*^-/y^ versus *Fmr1*^-/y^ *Raptor*^wt*/*f^ *CaMK2α*^Cre^, *p* = 0.0060, *q*_19_ = 5.00; B, PFC, *F*_2, 19_ = 14.52, *p* = 0.0001, *CaMK2α*^Cre^ versus *Fmr1*^-/y^, *p* = 0.0001, *q*_19_ = 7.486; *CaMK2α*^Cre^ versus *Fmr1*^-/y^ *Raptor*^wt*/*f^ *CaMK2α*^Cre^, *p* > 0.05; *Fmr1*^-/y^ versus *Fmr1*^-/y^ *Raptor*^wt*/*f^ *CaMK2α*^Cre^, *p* = 0.0033, *q*_19_ = 5.372, One-way ANOVA followed by Tukey’s multiple comparisons test) and p-eIF2α (C, hippocampus, *F*_2, 19_ = 14.28, *p* = 0.0002, *CaMK2α*^Cre^ versus *Fmr1*^-/y^, *p* = 0.0001, *q*_19_ = 7.549; *CaMK2α*^Cre^ versus *Fmr1*^-/y^ *Raptor*^wt*/*f^ *CaMK2α*^Cre^, *p* = 0.0129, *q*_19_ = 4.505; *Fmr1*^-/y^ versus *Fmr1*^-/y^ *Raptor*^wt*/*f^ *CaMK2α*^Cre^, *p* = 0.0304, *q*_19_ = 3.935; D, PFC, *F*_2, 19_ = 9.221, *p* = 0.0016, *CaMK2α*^Cre^ versus *Fmr1*^-/y^, *p* = 0.0012, *q*_19_ = 6.036; *CaMK2α*^Cre^ versus *Fmr1*^-/y^ *Raptor*^wt*/*f^ *CaMK2α*^Cre^, *p* > 0.05; *Fmr1*^-/y^ versus *Fmr1*^-/y^ *Raptor*^wt*/*f^ *CaMK2α*^Cre^, *p* = 0.0308, *q*_19_ = 3.927, One-way ANOVA followed by Tukey’s multiple comparisons test) expression in excitatory neurons. Each data point represents an individual animal. All data are presented as mean ± s.e.m. \**p* < 0.05, \*\**p* < 0.01, \*\*\**p* < 0.001, ns, not significant. Scale bars, 25 µm.

**Figure S6.**
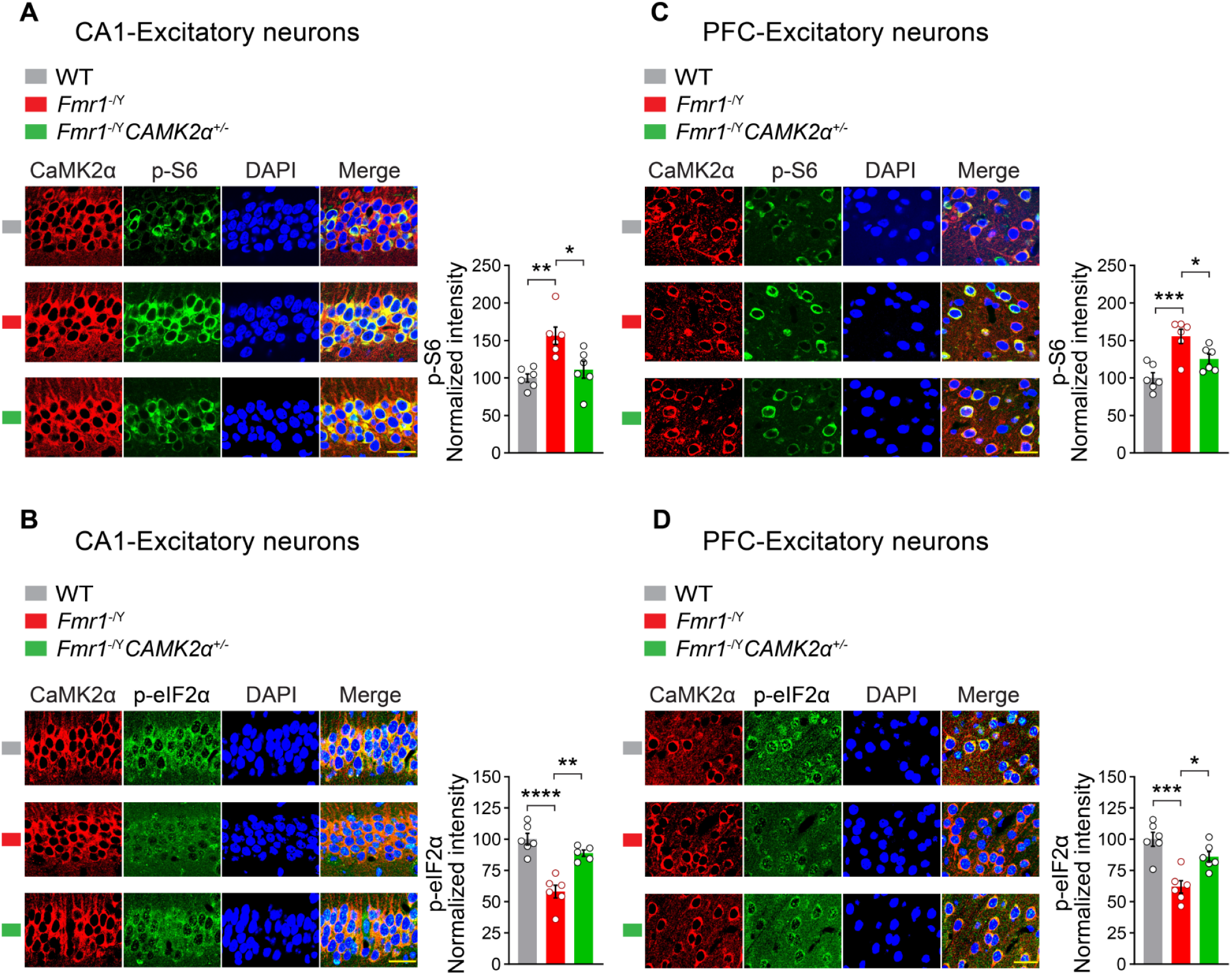
Deletion of one allele of *CaMK2α* in *Fmr1*^-/y^ mice rectifies elevated p-S6 and diminished p-eIF2α in excitatory neurons. (A-D) Representative images and quantification of p-S6 and p-eIF2α in the hippocampus and PFC of mice with heterozygous deletion of *CaMK2α* in *Fmr1*^-/y^ mice show a significant rescue in p-S6 (A, hippocampus, *F*_2, 15_ = 9.292, *p* = 0.0024, WT versus *Fmr1*^-/y^, *p* = 0.0027, *q*_15_ = 5.751, *Fmr1*^-/y^ versus *Fmr1*^-/y^ *CaMK2α*^+/-^, *p* = 0.0134, *q*_15_ = 4.628; B, PFC, *F*_2, 15_ = 12.34, *p* = 0.0007, WT versus *Fmr1*^-/y^, *p* = 0.0005, *q*_15_ = 7.016, *Fmr1*^-/y^ versus *Fmr1*^-/y^ *CaMK2α*^+/-^, *p* = 0.0403, *q*_15_ = 3.834) and p-eIF2α (C, hippocampus, *F*_2, 14_ = 24.80, *p* < 0.0001, WT versus *Fmr1*^-/y^, *p* < 0.0001, *q*_14_ = 9.658, *Fmr1*^-/y^ versus *Fmr1*^-/y^ *CaMK2α*^+/-^, *p* = 0.0008, *q*_14_ = 6.745; D, PFC, *F*_2, 15_ = 14.99, *p* = 0.0003, WT versus *Fmr1*^-/y^, *p* = 0.0002, *q*_15_ = 7.658, *Fmr1*^-/y^ versus *Fmr1*^-/y^ *CaMK2α*^+/-^, *p* = 0.0102, *q*_15_ = 4.820) in excitatory neurons. One-way ANOVA followed by Tukey’s multiple comparisons test was performed for all experiments and n = 5-6/group. All data are presented as mean ± s.e.m. \**p* < 0.05, \*\**p* < 0.01, \*\*\**p* < 0.001, \*\*\*\**p* < 0.0001, ns, not significant. Scale bars, 25 µm.

**Figure S7.**
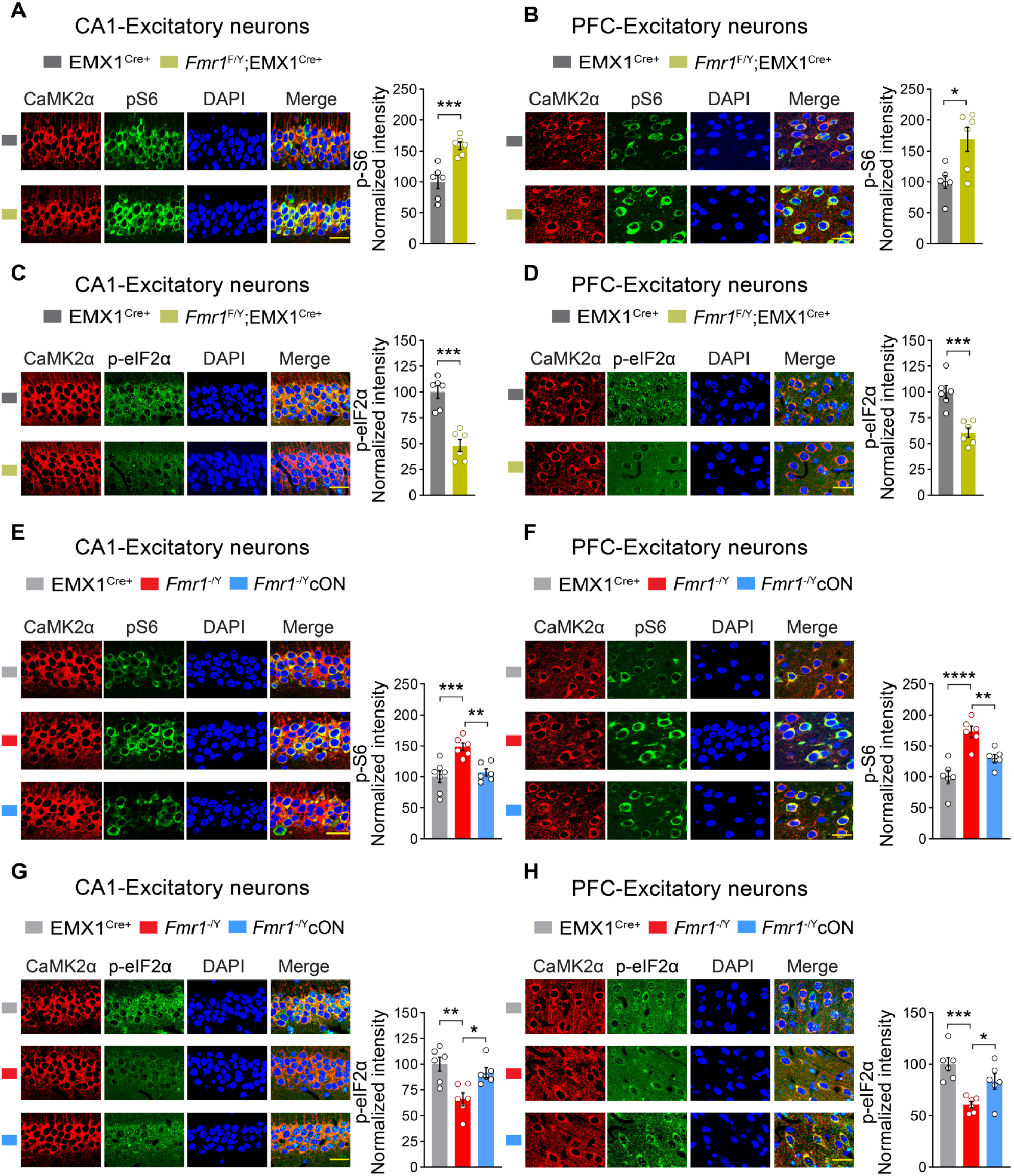
Autonomous effects of *Fmr1* ablation on p-S6 and p-eIF2α in excitatory neurons. (A-D) Deletion of the *Fmr1* gene in excitatory neurons causes an increase in p-S6 (A, hippocampus, *t* = 4.666, *p* = 0.0009; B, PFC, *t* = 3.124, *p* = 0.0108) and reduction in p-eIF2α (C, hippocampus, *t* = 6.053, *p* = 0.0001; D, PFC, *t* = 5.087, *p* = 0.0005) in *Fmr1*^-/y^ *EMX1*^Cre^ mice (n = 6) compared with WT animals (n = 6). Student’s t-test for all experiments. Excitatory neuron-specific expression of *Fmr1* gene in *Fmr1*^-/y^ mice normalizes the elevated p-S6 (E, hippocampus, *F*_2, 16_ = 11.70, *p* = 0.0007, *EMX1*^Cre^ versus *Fmr1*^-/y^, *p* = 0.0009, *q*_6_ = 6.447, *Fmr1*^-/y^ vs. *Fmr1*cON *EMX1*^Cre^, *p* = 0.0047, *q*_6_ = 5.303; F, PFC, *F*_2, 15_ = 18.21, *p* < 0.0001, *EMX1*^Cre^ versus *Fmr1*^-/y^, *p* < 0.0001, *q*_6_ = 8.492, *Fmr1*^-/y^ vs. *Fmr1*cON *EMX1*^Cre^, *p* = 0.0080, *q*_6_ = 4.993) and decreased p-eIF2α (G, hippocampus, *F*_2, 15_ = 9.084, *p* = 0.0026, *EMX1*^Cre^ versus *Fmr1*^-/y^, *p* = 0.0026, *q*_6_ = 5.778, *Fmr1*^-/y^ vs. *Fmr1*cON *EMX1*^Cre^, *p* = 0.0191, *q*_6_ = 4.376; H, PFC, *F2, 15* = 10.90, *p* = 0.0012, *EMX1*^Cre^ versus *Fmr1*^-/y^, *p* = 0.0009, *q*_6_ = 6.581, *Fmr1*^-/y^ vs. *Fmr1* cON *EMX1*^Cre^, *p* = 0.0450, *q*_6_ = 3.752). One-way ANOVA followed by Tukey’s multiple comparisons test was performed for all experiments (n = 6). All data are presented as mean ± s.e.m. \**p* < 0.05, \*\**p* < 0.01, \*\*\**p* < 0.001, ns, not significant. Scale bars, 25 µm.

**Figure S8.**
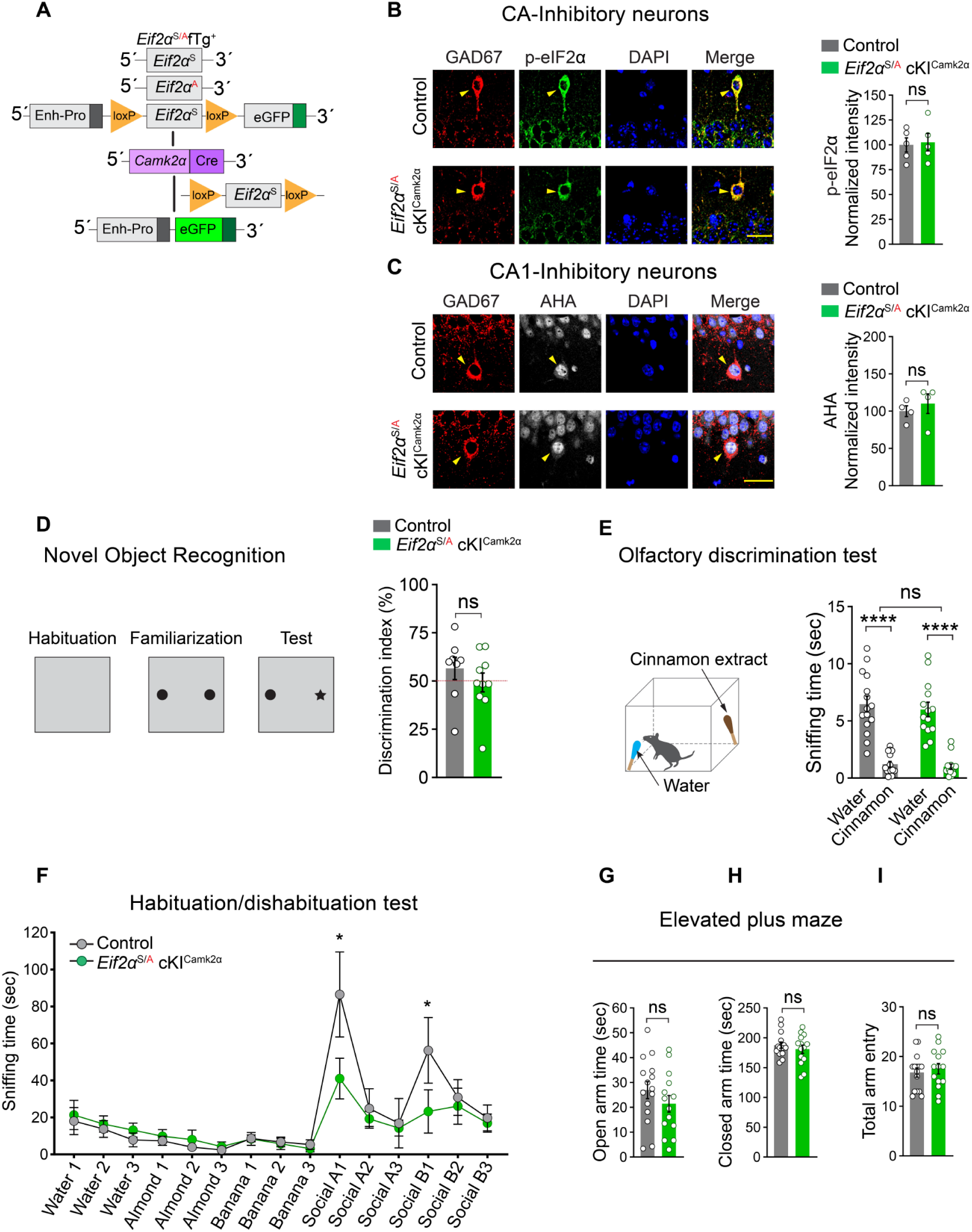
No changes in p-eIF2α and global protein synthesis in inhibitory neurons of *Eif2α*^S/A^ cKI^Camk2α^ mice. (A) A schematic illustration depicts the generation of *Eif2α*^S/A^ cKI^Camk2α^ mice (crossing the *Eif2α*^A/A^*fTg*^+^ mice with *Camk2*^Cre^ mice). Representative immunostaining and quantification of p-eIF2α (B) and AHA incorporation (C) in inhibitory neurons of hippocampal sections from *Eif2α*^S/A^ cKI^Camk2α^ mice (n = 4) show no differences in p-eIF2α levels (B, *t* = 0.242, *p* > 0.05, Student’s t-test) or AHA signal intensity (C, *t* = 0.679, *p* > 0.05, Student’s t-test) compared with control (*Camk2*^Cre^) mice (n = 4). Yellow arrows mark inhibitory neurons. (D) *Eif2α*^S/A^ cKI^Camk2α^ (n = 10) mice performed similarly to control animals in novel object recognition test (n = 8, *p* > 0.05, Student’s t-test). (E) *Eif2α*^S/A^ cKI^Camk2α^ mice exhibit intact olfaction. (E) Mice were allowed to freely sniff either the water or cinnamon extract-dipped swab. *Eif2α*^S/A^ cKI^Camk2α^ (n = 14) and control (n = 15) animals discriminated water over cinnamon extract, and no difference was found between the two groups (E, *F*_1, 54_ = 0.4543, *p* > 0.05, two-way ANOVA). (F) Olfactory habituation/dishabituation test. *Eif2α*^S/A^ cKI^Camk2α^ mice (n = 6) spent comparable amount of time to controls (n = 7) sniffing the non-social odors (water, almond extract, and banana extract), whereas *Eif2α*^S/A^ cKI^Camk2α^ mice show a significant impairment in sniffing the social odors (odor A and B, male urine) at the first trial (F, *F*_14, 154_ = 12.53, *p* < 0.0001, odor A-1, *t* = 4.657, *p* = 0.0183; odor B-1, *t* = 4.005, *p* = 0.0341, two way ANOVA followed by Bonferroni’s multiple comparisons test). (G-I) *Eif2α*^S/A^ cKI^Camk2α^ mice exhibit no anxiety-like behavior in an elevated plus maze test. *Eif2α*^S/A^ cKI^Camk2α^ mice (n = 14) spent a comparable amount of time to controls (n = 15) in open arm (G, *p* > 0.05) and closed arm (H, *p* > 0.05), and entered both arms as frequent as controls (I, *p* > 0.05). Student’s t-test was used for G-I. Each data point represents an individual animal. All data are presented as mean ± s.e.m. \**p* < 0.05, \*\*\*\**p* < 0.0001, and ns, not significant. Scale bars, 25 µm.

**Figure S9.**
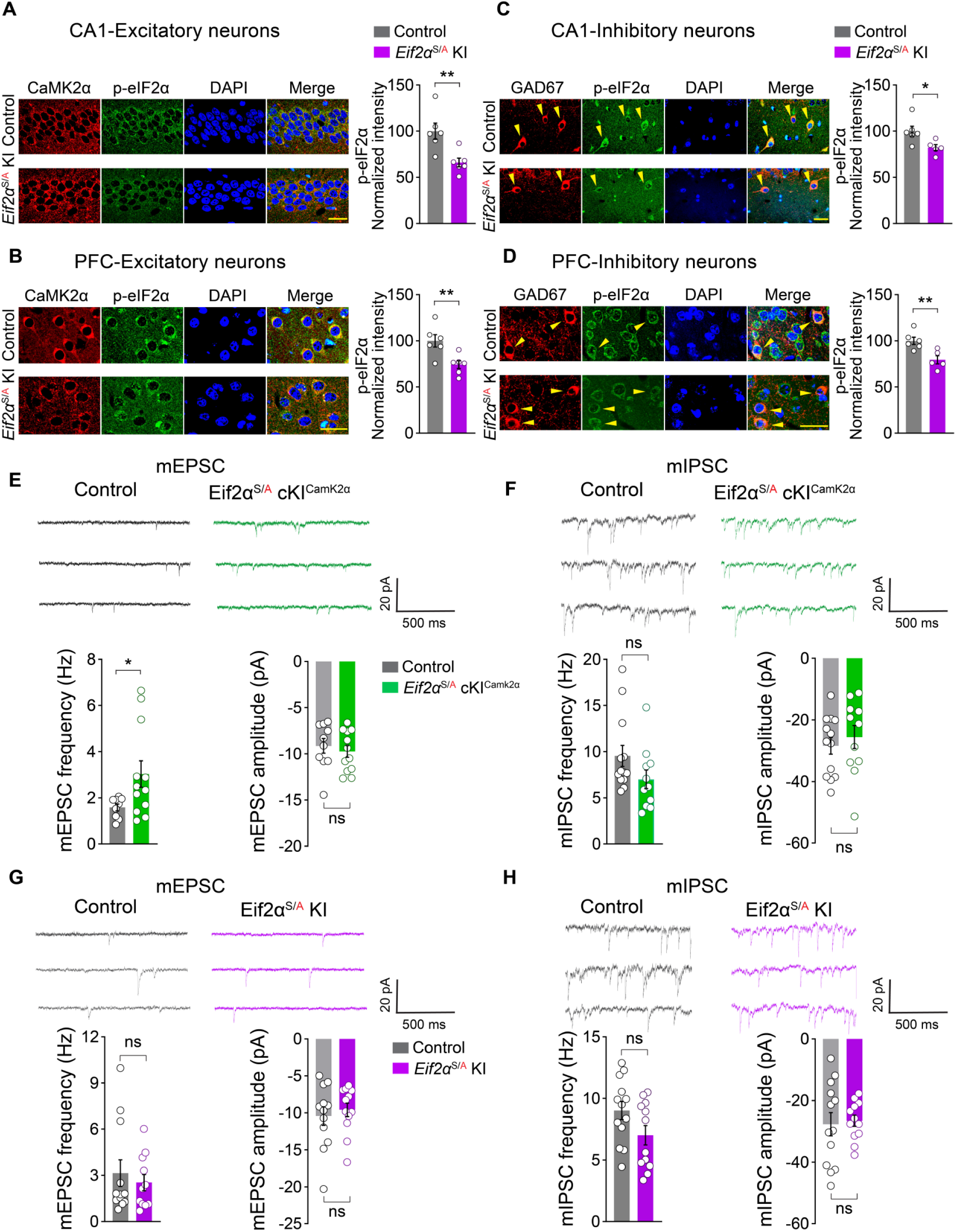
p-eIF2α is reduced in both excitatory and inhibitory neurons of *Eif2α*^S/A^ KI mice. (A-D) Representative and quantification of p-eIF2α expression shows a significant reduction in p-eIF2α levels in excitatory (A, hippocampus, *t* = 3.371, *p* = 0.0071; B, PFC, *t* = 3.211, *p* = 0.0093) and inhibitory neurons (C, hippocampus, *t* = 2.670, *p* = 0.0256 ; D, PFC, *t* = 3.494, *p* = 0.0068) in *Eif2α*^S/A^ KI mice (n = 5-6) in comparison to controls (n = 6). Student’s t-test was performed for all the experiments. Yellow arrows mark inhibitory neurons. Scale bars, 25 µm. (E-H) Representative traces and analysis of mEPSCs and mIPSCs recorded from CA1 pyramidal neurons in male *Eif2α*^S/A^ cKI^CamK2α^, *Eif2α*^S/A^ KI, and their controls. Frequency of mEPSCs in *Eif2α*^S/A^ cKI^CamK2α^ mice was significantly increased (E, left, *t* = 2.147, *p* = 0.0449). In *Eif2α*^S/A^ cKI^CamK2α^ mice, the amplitude of mEPSCs and frequency and amplitude of mIPSCs were not altered. In *Eif2α*^S/A^ mice, no change in amplitude and frequency of mEPSCs and mIPSCs were found (*p* > 0.05). Student’s t-test was performed for all the experiments. All data are presented as mean ± s.e.m. \**p* < 0.05, \*\**p* < 0.01, and ns, not significant.

**Figure S10.**
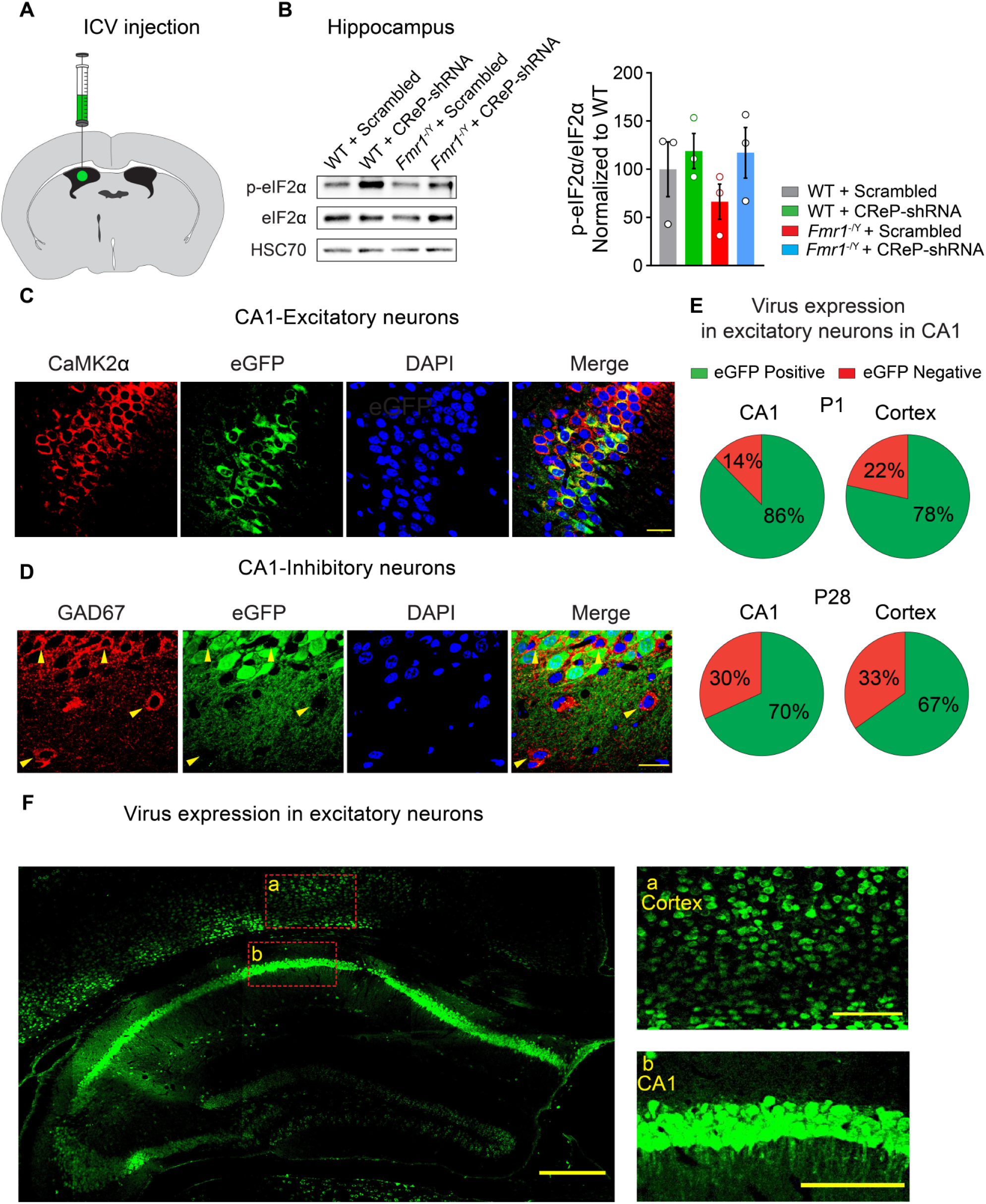
CReP-shRNA AAV is selectively expressed in Camk2α-positive neurons and upregulates p-eIF2α in *Fmr1*^-/y^ mice. (A) A schematic illustration shows the site of intracerebroventricular (i.c.v) AAV injection. (B) Immunoblots show that AAV9-Camk2α-GFP-*Ppp1r15b*-shRNAmir (CReP-shRNA) increased the level of p-eIF2α in the hippocampal lysates extracted from *Fmr1*^-/y^ mice (AAVs injected at P28, tissue collected at 8 weeks of age). (C, D) Representative immunostainings show that eGFP expression (driven by AAV) is restricted to excitatory neurons as eGFP (green) co-localizes with CaMK2α (C, red) but not with GAD67 (D, red). Scale bars, 25 µm. (E) CReP-shRNA AAV is expressed in significant fraction of excitatory neurons when AAVs are injected at P1 (top; CA1 86%, Cortex 78%) and P28 (bottom; CA1 70%, Cortex 67%). Yellow arrows mark inhibitory neurons. (F) A representative image shows CReP-shRNA AAV expression (eGFP) in hippocampus and cortex. (F, right) High resolution images of (a) cortex and (b) hippocampus. Scale bars; F, 300 µm; a and b,100 µm.

**Figure S11.**
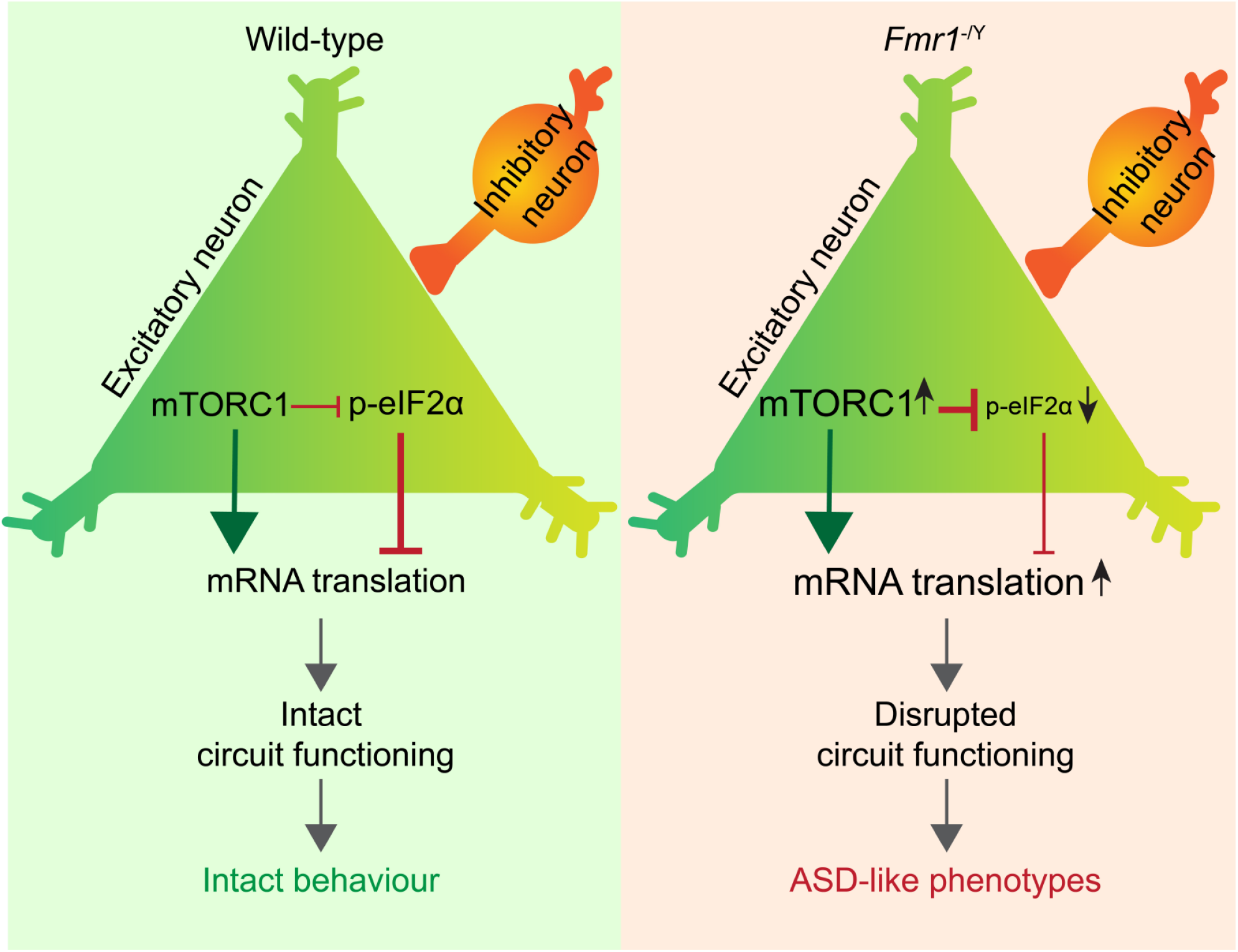
A model describing changes in the mTORC1 and ISR pathways in the mouse model of fragile X syndrome. mTORC1 activity is increased in excitatory but not inhibitory neurons of *Fmr1*^-/y^ mice as a result of elevated levels of CaMK2α in excitatory neurons. Increased mTORC1 causes eIF2α dephosphorylation that leads to increased protein synthesis in excitatory neurons. The resulting excitation/inhibition imbalance engenders neuronal circuit dysfunction and ASD-related behaviors.

## STAR Methods

### Animals

The *Fmr1*^−/y^ (Jackson Laboratories, stock #003025), general heterozygous knock-in (*Eif2α*^S/A^ KI),^43, 57^ *Eif2α*^S/A^ fTg^+^ floxed,^42^ *Fmr1*^f/f^,^56^ *Fmr1* conditional on (cON),^38^ *Raptor* floxed,^58^ *Camk2α*^+/-^ (Jackson Laboratories, stock #002362), *Camk2α*^Cre^ (Jackson Laboratories, stock #005359) and *EMX1*^Cre^ (Jackson Laboratories, stock #005628) mice were on the C57BL/6 background. To obtain the *Eif2α*^S/A^ fTg^+^; *Camk2α*^Cre+^ (*Eif2α*^S/A^ cKI^Camk2α^) mice, *Eif2α*^A/A^ fTg^+^ floxed mice^42^ were bred with *Eif2α*^S/S^ fTg^+^ *Camk2α*^Cre+^ mice. The heterozygous knock-in *Eif2α*^S/A^ fTg^+^; *Camk2α*^Cre+^ mice were then crossed with the same line to generate the homozygous mice *Eif2α*^A/A^ fTg^+^; *Camk2α*^Cre+^ (*Eif2α*^A/A^ cKI^Camk2α^). *Eif2α*^S/A^ fTg^+^ floxed, and *Camk2α*^Cre+^ mice were used as controls in behavioral experiments. No differences between these lines were observed, and the data were pooled and presented as “Control”. All experiments were conducted on adult male mice (8-week-old), except for electrophysiology and audiogenic seizure experiments in which young male mice (4–5-week-old) were used. All experiments were performed and analyzed by an experimenter blind to genotypes and treatments. Animals were housed in regular Plexiglas mouse cages with food and water available *ad libitum* and kept on a 12 h light/dark cycle (lights on at 7:00 AM). All procedures were compliant with the Canadian Council on Animal Care guidelines and approved by the McGill University’s Animal Care Committee.

### Stereotaxic surgery

Mice were deeply anesthetized (induced with 3% isoflurane and maintained on 1.5 % isoflurane) and placed in a stereotaxic frame (Kopf). The front teeth were placed in the incisor bar, and the head was secured using ear bars. The skull was exposed through a midline incision and drilled at the defined coordinates to microinject the AAV into the lateral ventricles (i.c.v.) (anterior/posterior (AP): -0.55 mm, medial/lateral (ML): 1.1 mm, and dorsal/ventral (DV): -2.2 mm) or CA1 hippocampal area (AP: -1.90 mm, ML: ±1.0 mm, and DV: -1.50 mm). Using a 10 µl Hamilton syringe connected to a 23-gauge needle and mounted on a perfusion pump, mice received 7 µl of AAV9-Camk2α-GFP-*Ppp1r15b*-shRNAmir (3.8 x 10^13^ GC/mL) or control AAV9-Camk2α-GFP-scrambled-shRNAmir (2.6 x 10^13^ GC/mL) via I.C.V. injection. The perfusion rate was set at 0.5 µl /min, and the needle was kept in place for additional 3 minutes before the withdrawal. Experiments were performed four weeks post-viral injection. For mGluR-LTD (Figure 6B-D) and audiogenic seizure (Figure 6I-K) studies, postnatal day 1 pups were injected i.c.v. and experiments were performed 4-5 weeks later (since mGluR-LTD and audiogenic seizure phenotypes in *Fmr1*^-/y^ mice are most robust at this age).^45, 59–61^ Specifically, pups received 2 µl of AAV9-Camk2α-GFP-*Ppp1r15b*-shRNAmir or AAV9-Camk2α-GFP-scrambled-shRNAmir i.c.v. bilaterally. Two-fifths of the hypothetical line from lambda to eyes was marked as the coordinate for the i.c.v. microinjection site. Then, the AAV was microinjected using a 5 µl Hamilton syringe connected to a 23-gauge needle. The needle was inserted 3 mm deep into the lateral ventricle.

### AAV9-shRNAmir cloning and preparation

The microRNA-adapted short hairpin RNAs (shRNAmir) packaged in adeno-associated virus (AAV) were prepared by Vector Biolabs. The validated sequence targeting mouse *Ppp1r15b* was: *GCTGTGAACTCAGAGACTTCTGCACGTTTTGGCCACTGACTGACGTGCAGAACTCTGAGTTC ACAG,* and the scrambled sequence used as a control was: GCGAGTCTCCACGCGCAGTACATTTTAGTGAAGCCACAGATGTAAAATGTACTGCGCGTGGAGACCTGC

### Western blotting

Mice were decapitated following deep anesthesia, and the brains were extracted and immediately placed in a dish containing ice-cold phosphate-buffered saline (PBS) to dissect the hippocampus, amygdala, and cortex. The extracted tissues were homogenized in a tissue homogenization buffer composed of 200 mM HEPES, 50 mM NaCl, 10% Glycerol, 1% Triton X-100, 1mM EDTA, 50 mM NaF, 2 mM Na3VO4, 25 mM β-glycerophosphate, and EDTA-free complete ULTRA tablets (Roche, Indianapolis, IN). Homogenized tissue was then centrifuged (14000 rpm for 15 min at 4 °C) to obtain the supernatant. Bradford protein assay was performed to measure protein concentration of the lysates. 30 µg of the lysates were loaded on a 12 % SDS-PAGE gel and ran at a constant current (0.03 A/per gel). The gel was then transferred to a nitrocellulose membrane overnight at a constant voltage (125 V). Next, the membrane was blocked for one h in 5% milk or BSA in TBS-T. The blocked membrane was incubated in primary antibodies (the list of antibodies is provided in the supplementary table 1) overnight at 4 °C. Primary antibody incubation was followed by three washes and incubation with HRP-conjugated secondary antibody (1:5000, Cat. No. NA931-1ML, Amersham) at room temperature. Then, the membrane was washed again, and the signal was enhanced using Enhanced Chemiluminescent (ECL) reagent and visualized using a ChemiDoc Imaging System (Bio-Rad). Antibody used in this study were: phospho-eIF2α (Ser51) (Cell Signaling Technology, Cat No. 3398), eIF2α (Cell Signaling Technology, Cat No. 9722), Anti-FMRP (Cell Signaling Technology, Cat No. 4317), GCN2 (phospho T899) (Abcam, Cat No. ab75836), GCN2 (Cell Signaling Technology, Cat NO. 3302), p-PERK (Thr 981), (Santa Cruz Biotechnology, Cat No. sc-32577), PERK (H-300), (Santa Cruz Biotechnology, Cat No. sc-13073), p-PKR (Thr 451) (Santa Cruz Biotechnology, Cat No. sc-101784), PKR (B-10) (Santa Cruz Biotechnology, Cat No. sc-6282), GAPDH (0411) (Santa Cruz Biotechnology, Cat No. sc-47724), GADD34 (Proteintech, Cat No. 10449-1-AP), CReP (PPP1R15B) (Proteintech, Cat No. 14634-1-AP), phospho-S6 ribosomal protein (Ser240/244) (Cell Signaling, Cat No. 5364), S6 ribosomal protein (5G10) (Cell Signaling Technology, Cat No. 2217), PP1C gamma antibody [EPR8934] (Abcame, Cat No. ab134947), and HSC 70 (K-19) (Santa Cruz Biotechnology, Cat No. sc-1059).

### Immunohistochemistry

Mice were transcardially perfused with 4% paraformaldehyde (PFA). The brains were postfixed in PFA 4% for 24 h following extraction. The fixed brains were sectioned using a vibratome to acquire 50-µm-thick sections. After washing with PBS, sections were blocked using 10% normal goat serum (NGS) and 0.5% Triton-X100 in PBS for two hours. Next, the blocked sections were incubated in primary antibody (provided in the supplementary table 1) diluted in PBS containing 2% NGS overnight. Sections were washed three times with PBS and incubated for one hour at room temperature in secondary antibody diluted in PBS. DAPI (1:5000) was added to the solution in the last wash. Rinsed sections were mounted on glass slides, imaged using a confocal microscope (Zeiss LSM 880), and scored using ImageJ (NIH). In all experiments, confocal imaging with 63X Oil objective (NA 1.4) was used, except for the 20X objective for eIF2α imaging. Two sections per mouse from each brain area were imaged (n = 4-6 mice/group). All images were captured using Z-stack mode with 15-20 optical sections/stack. For excitatory neurons, integrated density of p-eIF2α and p-S6 signal in the cytoplasm of 40 neurons per mouse (2 sections per mouse, 20 neurons per section) in the hippocampus, prefrontal cortex, and basolateral amygdala were quantified using ImageJ on maximum intensity projection images. The values from 40 neurons per mouse were averaged to obtain a single value per mouse. For inhibitory neurons, 16-20 neurons per mouse (8-10 neurons per section) were quantified as described for excitatory neurons. Measurements were restricted to cytoplasm using nuclear DAPI stain. Antibody used in this study were: CaMKII-α (6G9) (Cell Signaling Technology, Cat No. 50049), GAD67 (Sigma-Aldrich, Cat No. MAB5406), EIF2S1 (phospho S51) (Abcam, Cat No. ab32157), eIF2α (L57A5) (Cell Signaling Technology, Cat No. 2103), and phospho-S6 ribosomal protein (Ser240/244) (Cell Signaling Technology, Cat No. 5364).

### Marble burying test

After handling and habituation to the test environment, mice were individually placed on top of a 5 cm thick and leveled bedding with 18 shiny and clean marbles (distributed in 3 rows of 6 marbles) in a Plexiglas box (50 cm x 50 cm x 31 cm). Mice were allowed to explore and bury the marbles for 30 min. A marble was counted as buried if more than 2/3 of it were covered by bedding.

### Self-grooming test

Mice were placed in a clean mouse cage with fresh beddings and allowed to habituate and explore the cage for 20 minutes. Grooming time in the last 10 min was scored and analyzed. Total time of self-grooming was measured using a manual stopwatch in the recorded video through a rigged camera in front of the cage.

### Elevated plus maze

The handled mice were individually placed at the intersection of the four arms facing one of the closed arms and were allowed to explore the maze for 5 min while their motion path was filmed using a mounted camera over the maze. The maze comprises 4 Plexiglas arms (two closed and two open arms) raised 50 cm above the ground level with height: 30 cm, length: 50 cm, and width: 10 cm. The time spent in either arm and the total number of arms entries were scored.

### Reciprocal male-male social interaction test

After five-min habituation in a clean mouse cage with fresh bedding, mice were individually exposed to an unfamiliar age and sex-matched mouse (WT) for 10 min. The recorded video was then analyzed offline to measure the total time of interaction between the test mouse and the unfamiliar one.

### Three-chamber social interaction test

Three interconnected Plexiglas chambers (36 cm × 28 cm × 30 cm) divided by transparent walls were used. The experiment was conducted in three 10-min phases, including habituation, sociability, and novelty preferences. In the habituation phase, the test mice were individually placed in the middle chamber to freely explore all three chambers through doorways on the walls. In the sociability phase, the test mice were exposed to age, and sex-matched unfamiliar mice (stranger 1, C57BL/6J) which were confined in a small wire cage in one chamber, and the identical empty wire cage was located in the corresponding spot in the other chamber. In the novelty phase, the test mice were exposed to the same unfamiliar mouse (stranger 1) from the sociability phase, which is familiar now, and another age and sex-matched unfamiliar mouse (stranger 2, C57BL/6J) was placed in the empty wire cage. Once the doorways are opened, the test mice could explore both chambers containing stranger1 or stranger 2. All three phases were recorded using a mounted camera, and then the time spent sniffing of either empty or occupied wire cages, time spent in each chamber, and the total number of chamber entries were scored and analyzed. Any contact or sniffing when the test mouse approached as close as 1 cm of either empty or occupied wire cage was considered social interaction.

### Olfactory discrimination test

For assessing the primary olfactory function, mice were placed in a mouse cage with either water or cinnamon extract-soaked swabs. The swabs were placed on either side of the clean mouse cage. The subject mice were allowed to sniff and explore the swabs for 4 min. The time spent sniffing each swab was measured and analyzed.

### Novel Object Recognition

After three days of handling, mice were individually placed in an open-field box (60×60×30 cm) for 10 min to habituate to the box. In the next phase of the test known as the familiarization phase, mice were exposed to two identical objects for 5 min for two consecutive days. On the following day, mice were presented with two different objects (the original and a novel object) for 5 min. Novel object recognition was recorded and the total time of either sniffing or touching the object was measured by an investigator who was blind to genotypes. The discrimination index was calculated as DI = (novel object exploration time – familiar object exploration time/total exploration time) ×100.

### Habituation/dishabituation olfactory test

Following 30 min of habituation in the experimental room, 6–8-week-old male mice were exposed to cotton swabs containing five different odors in three consecutive trials (2 minutes) with 30 minutes inter trial intervals. Mice were exposed to the odors in the following order: water, almond extract, banana extract, social odor 1 and social odor 2. Social odors were collected from the dirty cages of stranger sex- and age-matched mice. Time spent sniffing each odor was recorded and analyzed.

### Audiogenic seizure test

Male mice (4-5-week-old) were individually placed in a soundproof Plexiglas cage and allowed to habituate for 2 min. Then a high-pitch siren (120 dB) was remotely turned on for 2 minutes. The number of wild running, tonic-clonic seizures, and status epilepticus/reparatory arrests were counted, and the percentage of incidents was analyzed. Note that status epilepticus is followed by an immediate respiratory arrest and death.

### Fluorescent non-canonical amino acid tagging (FUNCAT)

Mice were kept on a low-methionine diet for seven days. The next day, mice received azidohomoalanine (AHA) intraperitoneally (100 µg/gbw, i.p., Click-IT™ AHA (L-Azidohomoalanine), Cat No. C10102, Thermo Fisher Scientific). After 3 hours, mice were anesthetized and perfused transcardially with 4% PFA. The extracted brains were kept overnight in PFA 4% at room temperature 4°C for 24 h. Brains were then sectioned at 40 µm thickness. After washing, sections were blocked overnight at 4°C in blocking solution composed of 10% normal goat serum, 0.5% Triton-X100, and 5% sucrose in PBS. Sections were then “clicked” overnight in click buffer containing 200 μM triazole ligand, 400 μM TCEP, 2 μM fluorescent Alexa Fluor 647 alkyne (Alexa Fluor™ 647 Alkyne, Cat No. A10278, Thermo Fisher Scientific), and 200 μM CuSO4 in PBS. Sections were then washed and mounted on glass slides, imaged using a confocal microscope (Zeiss, LSM 880), and analyzed using ImageJ. FUNCAT images were acquired using an Airyscan mode on the Zeiss confocal microscope (LSM880) with 63X/1.40 Oil DIC f/ELYRA objective from two sections per mouse. Quantification was performed as described for immunohistochemistry. Forty excitatory and 16-20 inhibitory neurons were quantified per mouse.

### Pharmacological reagents

NV-5138 hydrochloride (Cat No. HY-114384B, MedChemExpress) was first dissolved in DMSO at 100 µg/µl. Then, it was further diluted in 5% Tween-80 and 40% PEG300 in saline (10.6 µg/µl) and administered via oral gavage at 160 mg/kg. Temsirolimus (CCI-779, Cat No. ab141999, Abcam) was first dissolved in DMSO at 50 µg/µl. Then, it was diluted in 5% Tween-80 and 5% PEG300 in saline (0.74 µg/µl) and injected at 7.5 mg/kg i.p. Anisomycin (Cat No. A9789, Sigma-Aldrich) was first dissolved in DMSO at 40 mM and brain slices (maintained in oxygenated ACSF) were incubated with anisomycin (40 µM) for 1h before AHA application.

### Field potential recordings

Following anesthesia induced by isoflurane, brains of 4-5-week-old male mice were rapidly extracted and dipped into an ice-cold sucrose-based cutting-solution (87 mM NaCl, 2.5 mM KCl, 1.25 mM NaH2PO4, 7 mM MgSO4, 0.5 mM CaCl2, 25 mM NaHCO3, 25 mM glucose, and 75 mM sucrose) and bubbled with 95% O2 and 5% CO2. Brains were sectioned using a vibratome to acquire transverse sections of 400 µm thickness. A surgical cut was performed to disconnect CA1 and CA3 regions of the hippocampus. Free-floating sections were allowed to recover in oxygenated and 32°C artificial cerebrospinal fluid (ACSF; 124 mM NaCl, 5 mM KCl, 1.25 mM NaH2PO4, 2 mM MgSO4, 2 mM CaCl2, 26 mM NaHCO3 and 10 mM glucose) for 2 h. One slice was then placed into the chamber and perfused with ACSF at 28 °C for an additional 30 min. Using a glass electrode (impedance; 2-4 MΩ) filled with ACSF, field excitatory postsynaptic potentials (fEPSP) were recorded from the CA1 stratum radiatum while the Schaffer collateral pathway was stimulated using a concentric bipolar tungsten stimulating electrode with 0.1 ms pulses at 0.033 Hz. The intensity was adjusted to evoke fEPSPs with 50% maximal amplitude. mGluR-dependent LTD was induced using group I mGluR agonist (S)-3,5-Dihydroxyphenylglycine (DHPG, Cat No. ab120007, Abcam) in ACSF perfusion (50 µM for 10 min). fEPSPs were recorded for one hour after induction of LTD. fEPSP slope between 10% and 90% of the maximal fEPSP amplitude was computed on Clampfit software. Fiber volley and population spikes were excluded from the analysis.

### Whole-cell recording

Whole-cell recording of synaptic currents: acute hippocampal slices (300 μm thickness) were prepared in ice-cold cutting solution containing (in mM): 75 sucrose, 87 NaCl, 2.5 NaH_2_PO_4_, 1.25 MgSO_4_, 0.5 CaCl_2_, 25 glucose, and 25 NaHCO_3_. Slices were transferred to artificial cerebrospinal fluid (ACSF) containing (in mM): 124 NaCl, 2.5 KCl, 1.25 NaH_2_PO_4_, 24 NaHCO_3_, 2 MgCl_2_, 2 CaCl_2_, and 12.5 glucose. Whole-cell recordings were obtained from CA1 pyramidal neurons using patch pipettes (borosilicate glass capillaries; 3–5 MΩ). To record miniature excitatory postsynaptic currents (mEPSCs), the intracellular solution contained (in mM): 130 CsMeSO_3_, 5 CsCl, 2 MgCl_2_, 5 diNa-phosphocreatine, 10 HEPES, 2 ATP-Tris, and 0.4 GTP-Tris (pH 7.3, 285 mOsmol). mEPSCs were recorded by voltage clamping the membrane at -70 mV in the presence of TTX 1 μM (0.5 μM; Alomone Labs) and gabazine (5 μM; Tocris Biosciences). To record miniature inhibitory postsynaptic currents (mIPSCs), the intracellular solution contained (in mM): 135 CsCl, 10 HEPES, 2 QX-314, 2 MgCl_2_, and 4 MgATP (pH 7.3; 285 mOsmol). mIPSCs were recorded by voltage clamping the membrane at -70 mV in the presence of TTX 1 μM, 6,7-dinitroquinoxaline-2,3-dione (DNQX 5 μM, Tocris Biosciences), and APV (50 μM, Tocris Biosciences). Recordings were obtained in voltage-clamp mode using a Multiclamp 700B amplifier (Molecular Devices). Recorded signals were digitized at 20 kHz and stored on a PC. Data acquisition and off-line analyses were performed using 1550B Digidata acquisition board, and pClamp 10 software (Molecular Devices). Data were included only if the holding current was stable. Detection threshold was set at 3 pA and 150–200 events were sampled per neuron.

### Analysis of 5**′** untranslated regions (5**′** UTR)

Analysis of UTRs of genes that showed changes in translation/ribosome association was carried out using a custom-scripted pipeline implemented in R. In brief, genes of interest were fed into the pipeline and UTR sequences retrieved from reference data (RefSeq; NCBI), according to user specifications (5′ and only longest UTR sequences). These sequences were then scanned for known UTR functional elements (motifs) using a stand-alone version of UTRscan.^62^ The data were then summarized to represent percentage (or number) of input genes harbouring one or more of the identified functional elements.

### Statistical analysis

GraphPad Prism 9 (GraphPad Prism Software Inc., USA) was used for statistical analysis. Data are shown as mean ± S.E.M., and the significance level was set at 0.05 or p < 0.05. A two-tailed unpaired Student’s t-test was performed to determine the differences between the two groups. Differences between multiple groups were determined using either one-way ANOVA or two-way ANOVA followed by either Tukey’s or Bonferroni’s post-tests. Fischer’s exact test (two-sided) was performed to analyze differences in the percent of seizure incidents and TRAP data. Statistical details of individual experiments are presented in figure legends. All experiments were scored by an experimenter blind to genotypes and treatments. Data points in all graphs represent the number of animals.

## Materials and Correspondence

Correspondence and requests for materials should be addressed to Nahum Sonenberg, Christos G. Gkogkas, and Arkady Khoutorsky. All data are available in the main text or supplementary materials.

